# Targeting the ubiquitin-proteasome system in a pancreatic cancer subtype with hyperactive MYC

**DOI:** 10.1101/2020.07.01.182162

**Authors:** K Lankes, Z Hassan, MJ Doffo, C Schneeweis, S Lier, R Öllinger, R Rad, OH Krämer, U Keller, D Saur, M. Reichert, G Schneider, M Wirth

## Abstract

**Purpose:** The myelocytomatosis oncogene (MYC) is an important driver in a subtype of pancreatic ductal adenocarcinoma (PDAC). However, MYC remains a challenging therapeutic target, therefore identifying druggable synthetic lethal interactions in MYC-active PDAC may lead to novel precise therapies.

**Methods:** Cluster analysis using direct MYC target genes was used to identify PDAC with active MYC. We profiled the transcriptome of established human cell lines, murine primary PDAC cell lines and also accessed public available repositories for transcriptomic profiling. Networks active in MYC hyperactive subtypes were analyzed by gene set enrichment analysis. An unbiased pharmacological drug screen with FDA-approved anti-cancer drugs was conducted to define MYC-associated vulnerabilities, which were validated by analysis of drug response repositories and genetic gain- and loss-of-function experiments.

**Results:** In an unbiased pharmacological drug screen with FDA-approved anti-cancer drugs we detected that the proteasome inhibitor bortezomib triggers a MYC-associated vulnerability. By integrating publicly available data sets we found the unfolded protein response as a signature connected to MYC. Furthermore, the increased sensitivity of MYC hyperactive PDACs to bortezomib was validated in genetically modified PDAC cells.

**Conclusions:** In sum, we provide evidence that perturbing the ubiquitin proteasome system might be an option to target MYC hyperactive PDAC cells and our data provide the rationale to further develop precise targeting of the ubiquitin-proteasome system as a subtype-specific therapeutic approach.

## Introduction

Pancreatic ductal adenocarcinoma (PDAC) is estimated to become the second leading cause of cancer-related death. In contrast to other solid tumors its prognosis still remains extremely poor [1]. The disease is characterized by a profound intertumoral heterogeneity [2]. Based on various technologies including mRNA sequencing, metabolite profiling, or exon sequencing, distinct molecular subtypes of PDAC associated with different prognosis, biology, and therapeutic responses have been described [2–13]. These data suggest the development of biomarker-driven therapeutic concepts as a promising approach to improve the outcome of the disease. Signatures predicting sensitivity towards the current standard of care chemotherapies are under development [12].

Whole-exome sequencing of micro-dissected PDAC specimen revealed that amplification of MYC (c-MYC) is the only copy number variation (CNV) associated with lower survival rates [14]. These data demonstrate that MYC drives an aggressive subtype of the disease and consistently MYC activity was found to be enriched in the squamous/basal-like/glycolytic subtype [7, 13]. MYC is involved in a variety of biological processes in cancer cells [15]. It is a prominent oncogene acting in concert with mutated *KRAS* in PDAC [16–18]. The transcription factor MYC is an intrinsically disordered protein. Although progress has been made to target MYC [19, 20], it remains a challenge. A strategy to target “undruggables” is to exploit specific cellular dependencies associated with the activity of these proteins [21]. Several unbiased genetic screens for synthetic lethal interactions have demonstrated that the MYC protein family confers targetable dependencies [22–26], pointing to a way to define precise therapies. Consistently, MYC has been connected to the increased sensitivity of bromodomain and extra terminal motif (BET) inhibitors [27, 28] and inhibitors of the small-ubiquitin-like modifier (SUMO) pathway in the context of PDAC [29].

To find MYC-associated vulnerabilities, we conducted a limited drug screen and found a connection of MYC to the unfolded protein response (UPR) and an increased sensitivity towards proteasome inhibitors.

## Results

### Drug screening of FDA-approved anti-cancer drugs identifies vulnerabilities in MYC^high^ human PDAC cells

To identify vulnerabilities in PDACs with an increased MYC activity, we performed an unbiased pharmacological drug screening experiment. To cover the identified MYC^high^ and MYC^low^ subgroups, we used four representative PDAC cell lines with low and high MYC expression (Fig. 1A). Except for PSN1, the doubling time of the cell lines were in similar ranges (Table S1). Gene set enrichment analysis (GSEA) with the GeneTrail2 1.6 web service [30] demonstrated the activation of the MYC network in cells with higher expression of the protein (Fig. 1B). A novel gene set of direct MYC target genes defined by Muhar et al. [31] showed the strongest enrichment in the MYC^high^ cell lines (Fig. 1B). We used the described models for a drug screening experiments with a set of 129 FDA-approved anticancer drugs, which is outlined in figure 2C. Hits were determined as a two-fold difference in the responsiveness of the MYC^high^ models. Among the ten candidates, we identified drugs from different classes, such as HDAC inhibitors, DNA antimetabolites, proteasome inhibitors, topoisomerase inhibitors, and others (Fig. 1D and Table S2).

**Figure 1.**
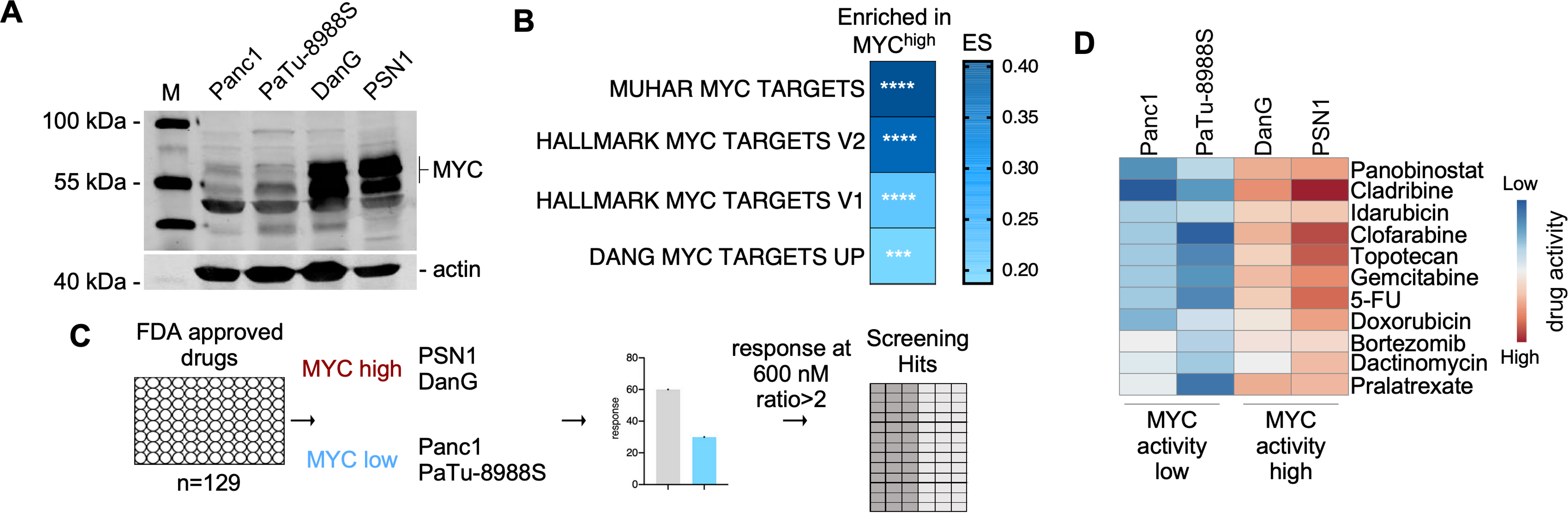
Drug screening in human PDAC cells with diverse MYC activity. **A)** MYC protein expression analysis of the four indicated PDAC cell lines was determined by western blotting. ACTB (beta-actin) served as a loading control. **B)** GSEA by GeneTrail2 1.6 web service demonstrates enrichment of the depicted MYC signatures in the MYC^high^ lines. Color coded enrichment score (ES) is depicted. *** adjusted p values<0.001; **** adjusted p values<0.0001. **C)** Strategy for drug screening experiments with of n=129 FDA-approved anticancer drugs. Cells were treated for 72h (two doubling times) with 600nM of each compound. Hits were determined as a two-fold difference in responsiveness. **D)** Top ten hits from the drug screening of 129 FDA-approved compounds depicted as a variance scaled heatmap.

### Validation experiments confirm drug screening results

To validate the single dose drug-screening experiment, we again examined the top 10 hits of our screening experiment using ten different doses and determined the dose-response curves. In addition to the used screening platform, we included a further PDAC line with low MYC protein expression (HPAC) and a PDAC line with intermediate/high MYC expression (MiaPaCa2). As shown in figure 2A, MYC protein expression is significantly different in the analyzed cell lines. Most dose response curves were all left-shifted in the MYC^high^ models (Fig. 2B). Despite the low MYC expression in HPAC cells, these cells cluster into the MYC^high^ high group and show increased sensitivity, which could be explained by expression of functional wild type p53. Although the mean area under the dose response curves (AUC) is lower for all screening hits in the MYC^high^ models, a high variance was detected (Fig. 2B). Such observations point to the need for large cell line panels to ultimate validate screening hits. Therefore, to further substantiate the screening hits, we accessed the discriminant regulon expression analysis (DoRothEA) database [32]. This database links the transcriptional activity of 127 transcription factors to drug sensitivity. We accessed the data for MYC and found significant overlaps of six drug classes with our screening experiment, which points to the robustness of the screen (Fig. 2C).

**Figure 2.**
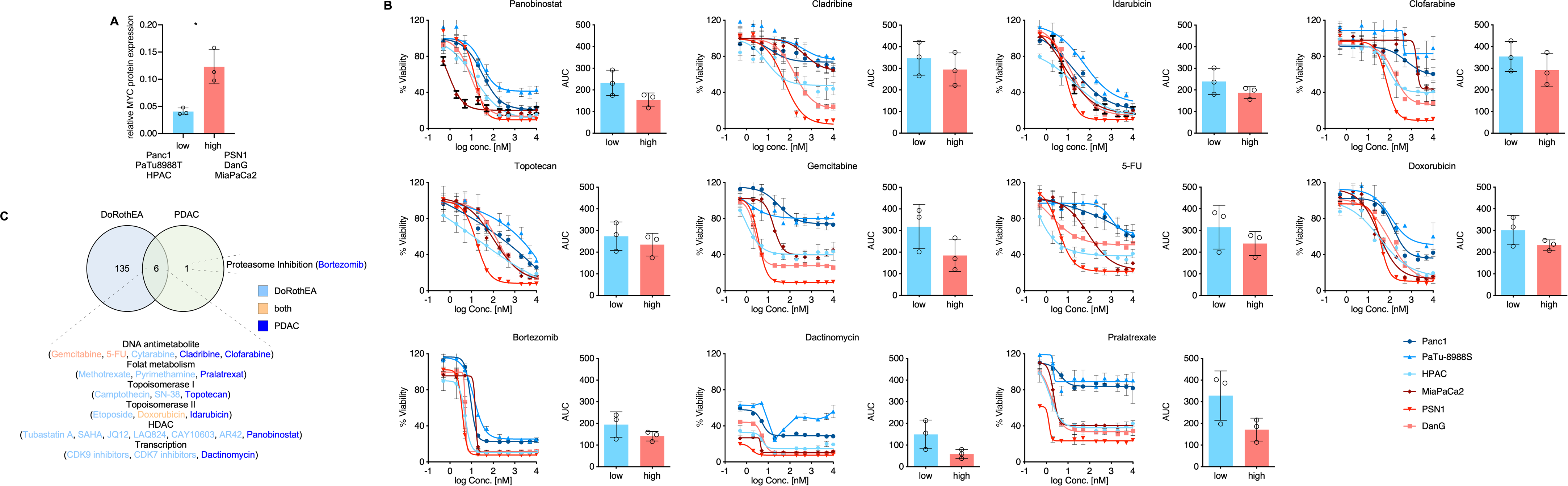
Validation of the drug screening experiment. **A)** Quantification of MYC expression of the indicated cell lines. In three independent lysates the MYC expression was determined and shown is the mean MYC expression per cell lines. *p value of an unpaired t-test <0.05. **B)** Viability for multi-dose treatment of MYC^high^ and MYC^low^ cells of displayed compounds. Cells were treated for 72h and viability was measured by MTT. Each dose-response experiment was conducted in three independent biological experiments conducted as technical triplicates. The area under the dose response curves (AUC) in both groups is depicted for each drug. **C)** Venn-diagram of data from the DoRothEA (Discriminant Regulon Expression Analysis) database and our drug screening hits. Significant (FDRq <0.05) drug-MYC interactions of the DoRothEA database were compared to the hits of our experimental drug screening experiment. Drugs hits were summarized into drug classes.

### MYC-associated pathways in human PDAC

To prioritize the hits of the screen, we accessed human PDAC mRNA expression datasets to observe potential connections of MYC-associated pathways to the screening hits. We used a PDAC data set from The Cancer Genome Atlas (TCGA) [33], which was curated according to Peran and colleagues (n=150) [34]. In addition, we used the International Cancer Gene Consortium (ICGC) dataset [7], in which acinar cell carcinomas and intraductal papillary mucinous neoplasms were excluded (n=81). We clustered both datasets according to the HALLMARK_MYC_TARGET_GENES_V1, the HALLMARK_MYC_TARGET_GENES_V2, and the MUHAR_MYC_TARGETS, as exemplified for the ICGC dataset in figure 3A. 11 cancers (approximately 10%) were defined as MYC hyperactivated by all three MYC signatures (Fig. 3A and 3B). These cancers showed a reduced survival (SFig. 1A) and a clear connection to the squamous subtype of the disease (Fig. 3A and SFig. 1B). Using the same approach for the TCGA dataset, 16 cancers (approximately 10%) were defined to be MYC hyperactivated by all three used MYC signatures (SFig. 1C). Although survival of MYC hyperactivated cancers was not reduced in this dataset (SFig. 1D), a connection to the basal-like cancer was again observed (SFig. 1E), which confirms the documented connection of MYC to this subtype of PDAC [7, 35]. To define MYC-associated pathways in the commonly MYC hyperactivated subtype, we performed a gene set enrichment analysis. 603 signatures were consistently enriched in both analyzed dataset when HALLMARKS-, KEGG-, GO-Term-, and REACTOME-signatures were accessed via the MSigDB (Fig. 3C). Figure 3D shows HALLMARK- and KEGG-signatures linked to MYC in both datasets. To corroborate the direct connection of such pathways, we used a MYC estrogen receptor fusion protein (MYC^ER^) of IMIM-PC1 cells, which are characterized by low MYC protein abundance [29]. Here, MYC-activated signatures show an overlap to the HALLMARK- and KEGG-signatures detected in the analysis of the ICGC and TGCA datasets, pointing to direct effects of MYC (SFig. 1F). Investigating the MYC-connected signatures, we detected a prominent proportion of ribosomal and translational signatures in both datasets, which is well in line with a recent analysis of the TCGA dataset demonstrating that translation is a key process linked to MYC in PDAC [36]. Consistent with increased translational activity, we detected signatures of the unfolded protein response (UPR) and UPR-activated signaling, including Proteinkinase RNA-activated-like ER Kinase (PERK) and Activating Transcription Factor 4 (ATF4) signatures in both investigated human PDAC datasets (Fig. 3E) [37].

**Figure 3.**
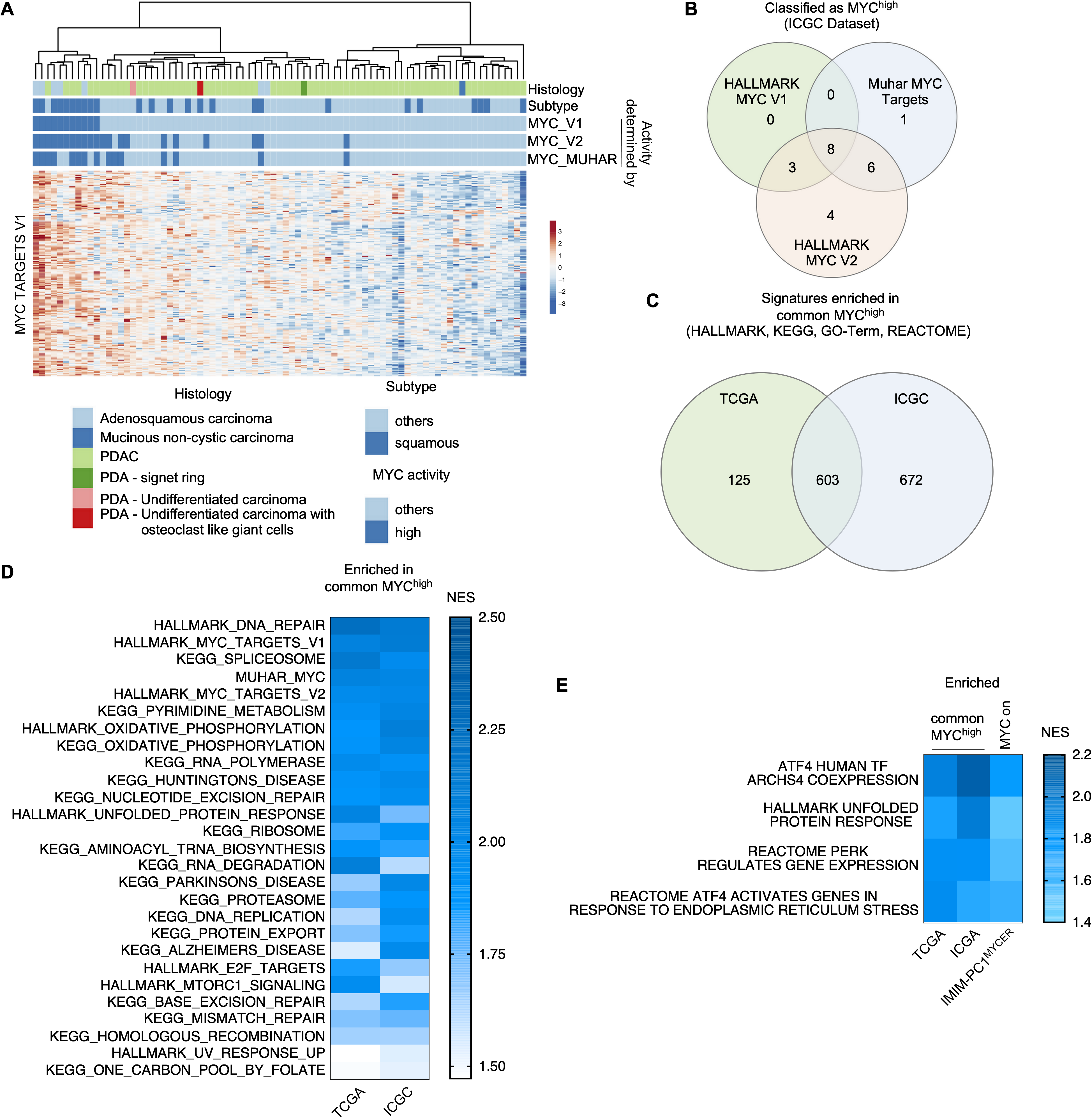
Pathways enriched in human common MYC^high^ PDACs. **A)** Clustering of the ICGC PDAC mRNA expression dataset according to the genes of the HALLMARK_MYC_TARGETS_V1 signature. Color coded information of the Histology, the Subtype and the MYC activity state determined by clustering of the HALLMARK_MYC_TARGETS_V1, the HALLMARK_MYC_TARGETS_V2 and the MUHAR MYC TARGETS [31] are depicted. **B)** Venn diagram of PDAC identified as MYC^high^ by clustering of the genes of the depicted signatures in the ICGC dataset. 8 PDACs were identified as common MYC^high^ PDACs. **C)** Common MYC^high^ PDACs of the TCGA and the ICGC dataset were analyzed by GSEA using the HALLMARK, the KEGG, the RREACTOME, and the GO-TERM signatures of the MSigDB with a FDR q value threshold of <0.25. The Venn diagram depicts 603 signatures enriched in common MYC^high^ PDACs of both datasets. **D)** NES visualized by a heatmap of the HALLMARK and the KEGG signatures enriched in common MYC^high^ PDACs of both datasets. **E)** NES visualized by a heatmap of gene signatures of the UPR and UPR-associated pathways enriched in common MYC^high^ PDACs of both datasets. As a control, IMIM-PC1^MYCER^ cells were used. Shown is the NES of the same signatures enriched in 4-OHT treated (MYC on) cells. For all depicted signatures: FDR q<0.05.

### MYC and sensitivity towards perturbants of protein homeostasis

The observation that MYC activity is connected to the UPR (Fig. 3D and 3E) and our recent demonstration that MYC is mechanistically involved in the induction of apoptosis in response to proteasome inhibition in PDAC cells [38], prompted us to investigate the bortezomib screening hit in greater detail. First, we used the dependency map (DepMap) portal to access Bortezomib sensitivity data for PDAC cell lines using data from the PRISM repurposing primary screen [39], the GDSC2 screen [40], and the CTD^2 screen [41]. We determined Bortezomib sensitive PDAC cell lines and analyzed them by a gene set enrichment analysis. Consistently, in all three datasets, we observed an enrichment of MYC signatures in the Bortezomib sensitive phenotype (Fig. 4A). To validate a connection of the MYC network to increased sensitivity towards proteasome inhibitors across species, we performed multi-dose drug screenings with the proteasome inhibitors Marizomib and Bortezomib in 38 well-characterized murine *Kras*^*G12D*^-driven PDAC cell lines [42] (Fig. 4B). The GI_50_ values of both inhibitors showed a significant correlation (Fig. 4B). We used RNA-seq data [42] of these murine PDAC lines and investigated enrichment of MYC signatures in Bortezomib sensitive, Marizomib sensitive, and lines sensitive to both proteasome inhibitors. We detected enrichment of the MUHAR MYC TARGETS and the HALLMARK MYC TARGETS V2 signature enriched in all proteasome inhibitor sensitive phenotypes (Fig. 4C). To test whether sensitivity of perturbants of the protein homeostasis are commonly connected to increased MYC activity, we analyzed the HSP90 inhibitors Ganetesip and NMS-E973, and the valosin-containing protein (VCP)/p97 inhibitor NMS-873. Such inhibitors are able to induce ER stress and the UPR [43–46]. Two additional proteasome inhibitors, Oprozomib and Ixazomib, were included as controls. To investigate the connection of HSP90 inhibitors and p97 inhibitors to MYC, we used again the data of the PRISM repurposing primary screen [39]. In GSEA, HSP90- and p97-inhibitors sensitive PDAC cell lines enrich for MYC-signatures and an unfolded protein response signature (SFig. 1B). The same was again observed for Oprozomip and Ixazomib sensitive lines (SFig. 1D). These data support the conclusion that MYC hyperactivated PDACs are more sensitive to perturbants of the protein homeostasis.

**Figure 4.**
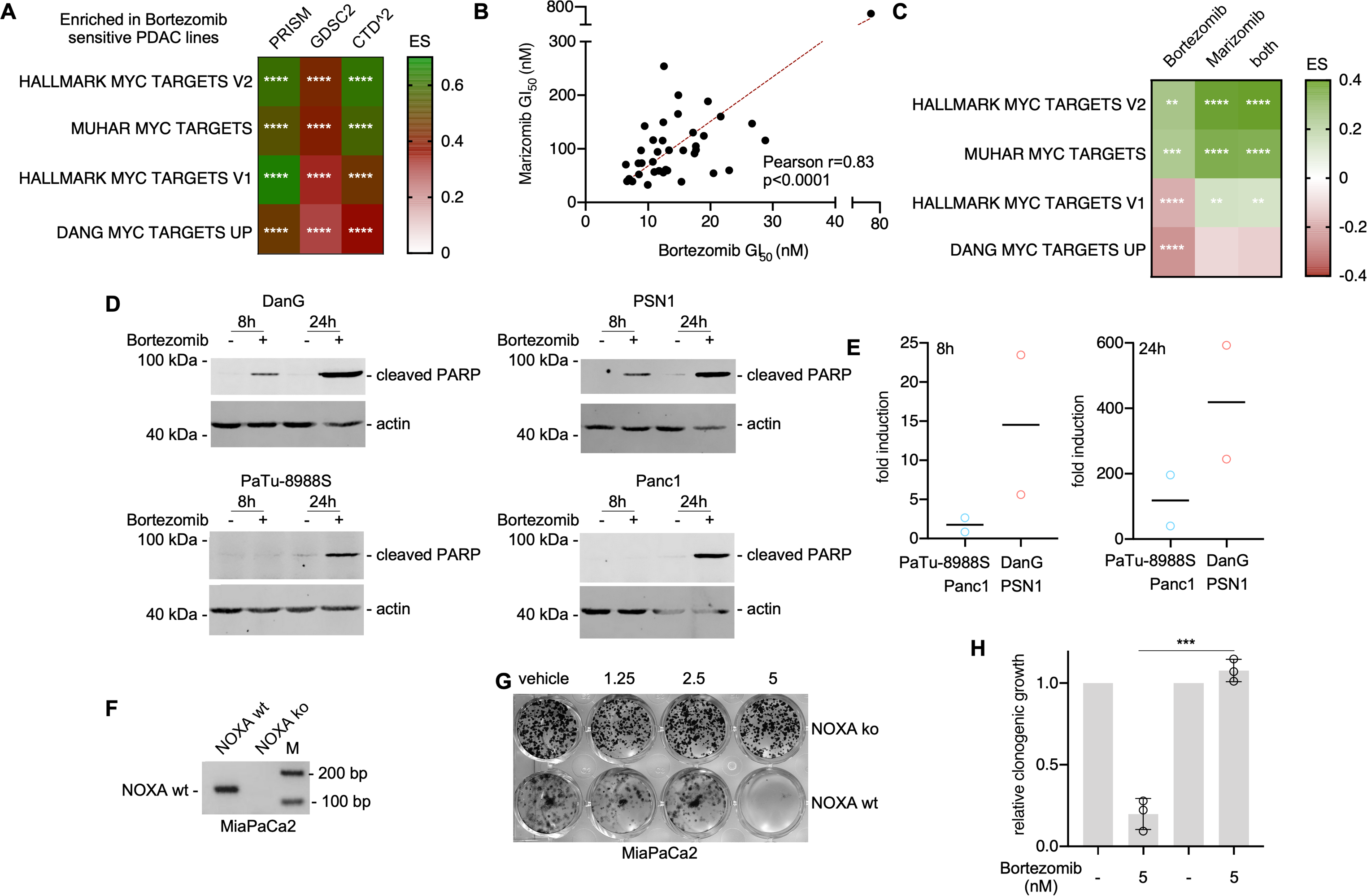
MYC primes for proteasome inhibitor induced apoptosis. **A)** Bortezomib sensitivities of human PDAC cell lines from the PRISM repurposing primary screen (19Q3) [39], the GDSC2 screen (AUC) [40], and the CTD^2 (AUC) screen [41] were divided into quartiles and lines for the most sensitive quartile were compared to the remaining cell lines of the complete CCLE PDAC dataset with a gene set enrichment analysis using the GeneTrail2 1.6 web service. The enrichment score was color coded. **** adjust. p-value<0.0001. **B)** Growth inhibitory 50% concentration of n=38 murine PDAC cell lines for Bortezomib and Marizomib was determined (72h of treatment, seven-point dilution, MTT assay, non-linear regression, n=3 independent biological replicates as technical triplicates). Depicted is the Pearson correlation coefficient and the linear regression (in red). **C)** Bortezomib and Marizomib GI50 values were divided into quartiles and lines from the most sensitive quartile were compared to the remaining cell lines by GSEA. In addition, the lines belonging to the Bortezomib as well as the Marizomib most sensitive quartile were compared to the rest of the lines by GSEA. GSEA was conducted by the GeneTrail2 1.6 web service. Color coded enrichment score is depicted. **adj. p-value<0.01, ***adj. p-value<0.001, ****adj. p-value<0.0001. **D)** Western blot analysis to determine expression of MYC, cleaved PARP and ACTB (loading control), 8 and 24 hours after treatment with 50nM Bortezomib or DMSO (vehicle control) (n=3). E) The cleaved PARP band was quantified in three independent expriments and the mean fold induction of cleaved PARP expression in MYC^low^ and MYC^high^ subtypes is depicted. **F)** Determination of CRISPR/Cas9 mediated knockout of the NOXA gene in MiaPaCa2 cells by PCR. A product size of 137 bp indicates the wild type allele, while no product indicates NOXA knockout a cells as described in MM section. **G)** Clonogenic growth assay of Bortezomib treated MiaPaCa-2 NOXA knockouts and wildtype cells with the indicated concentrations. One representative experiment out of three is depicted. **H)** Quantification of three independent clonogenic growth assays according to G). *p value of an unpaired t-test<0.001.

### Human PDAC cells with active MYC are primed for Bortezomib-induced apoptosis

Previously, we described that Bortezomib-induced apoptosis of PDAC cell lines is mediated by MYC-dependent activation of pro-death BCL2 family members, including NOXA (PMAIP1) [38]. To corroborate augmented apoptosis induction as the underlying principle for the increased sensitivity towards proteasome inhibition in MYC^high^ PDAC lines, we monitored cleavage of the caspase substrate PARP over time. Only in MYC^high^ lines, a significant cleavage of PARP was observed eight hours after the treatment (Fig. 4D). 24 hours after the treatment, caspases were also activated in MYC^low^ cell lines (Fig. 4D). Nevertheless, bortezomib induced PARP cleavage was always higher in MYC^high^ lines (Fig. 4E). Since the BH3-only pro-apoptotic BCL2 family member NOXA was recently described to contribute to bortezomib induced apoptosis in PDAC cell lines [38], we induced a CRISPR-Cas9 mediated knockout of the *NOXA* gene in MiaPaCa2 cells (Fig. 4F). The therapeutic response towards Bortezomib is distinctly reduced in *NOXA*-deficient MiaPaCa2 cells (Fig. 4G and 4H), demonstrating the relevance of the gene for the Bortezomib-induced apoptosis.

To analyze the direct contribution of MYC to the proteasome inhibitor sensitivity, we used the dual recombination system [47] with floxed *Myc* alleles [48] to generate a genetic loss of function PDAC model (Fig. 5A). Activation of a *Cre*^*ERT2*^ fusion by the addition of 4-hydroxytamoxifen (4-OHT) in this murine PDAC cell lines deleted the floxed *Myc* alleles and reduces MYC protein expression to approximately 30% compared to controls (Fig. 5B and 5C). It is important to note that we were not able to generate a complete MYC knock-out, due to the profound counter selection of recombination escapers, which underscores the importance of MYC as a target in PDAC. Nevertheless, the MYC-reduced population was less Bortezomib sensitive (Fig. 5D). In addition, we used a conditional gain-of-function model relying on a MYC estrogen receptor fusion (MYC^ER^). We transduced a murine PDAC cell line with low MYC expression. Upon treatment with 4-hydoxytamoxifen (4-OHT) the MYC targets *Odc1* and *Cad* were induced and endogenous *Myc* was repressed by its negative autoregulation (Fig. 5E). Seeding the cells in 4-OHT for 24 hours followed by a three-day treatment period with Bortezomib, did not change the sensitivity to the proteasome inhibitor (Fig. 5F). MYC-amplified PSN1 cells were included as a Bortezomib sensitive control. Considering the time needed to adapt the system to MYC, we followed two strategies. First, pre-treating the cells with 4-OHT for 24 hours followed by a six-day treatment period with Bortezomib, demonstrated increased sensitivity in the MYC “on” state (Fig. 5G). Second, pretreating the cells with 4-OHT for 96 hours followed by 72 hours of Bortezomib treatment also sensitized the cells to Bortezomib (Fig. 5H). Therefore, gain- and loss-of-function models support the note that MYC modulates the proteasome inhibitor sensitivity of PDAC cells.

**Figure 5.**
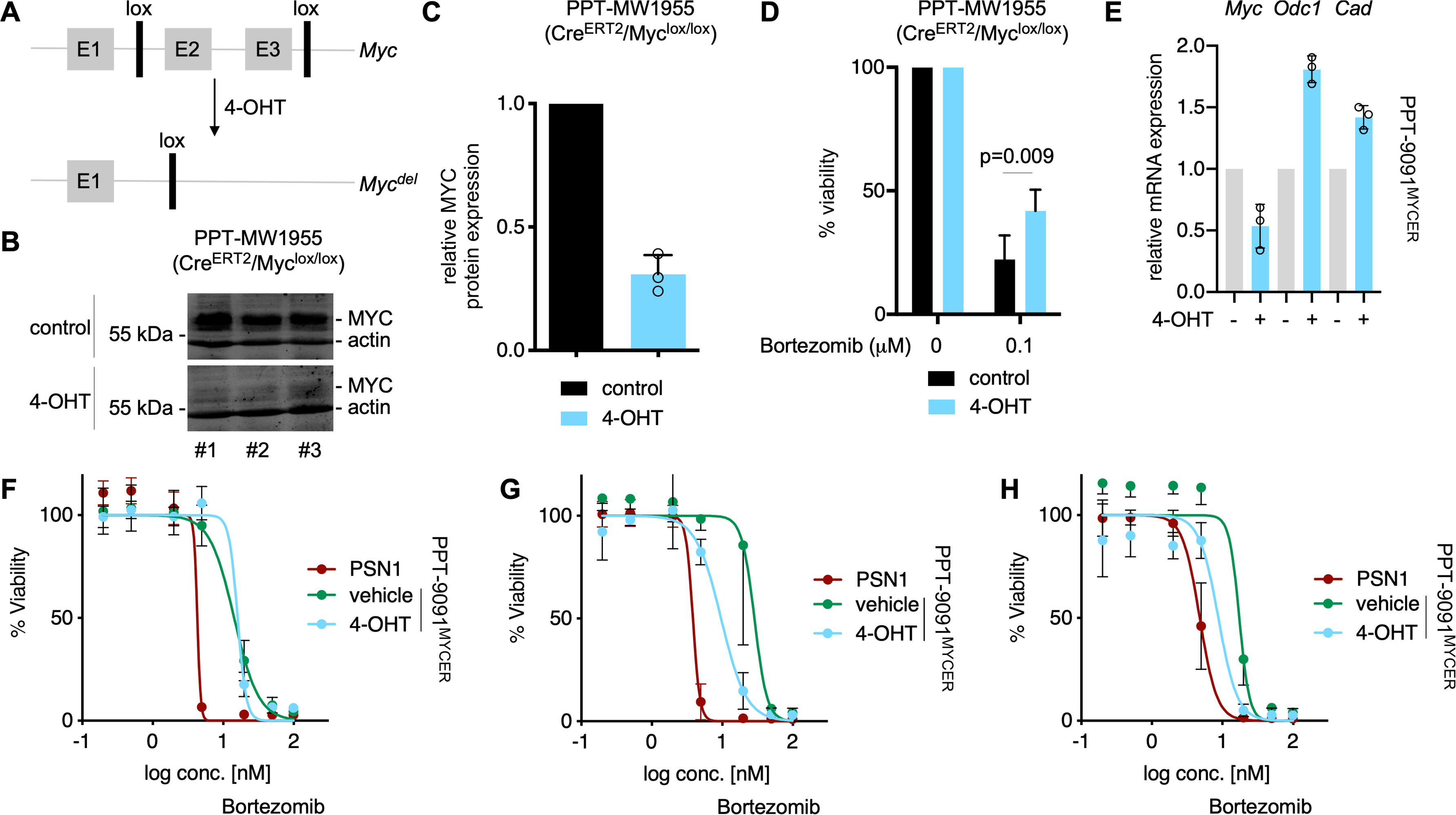
Proteasome inhibitor sensitivity and MYC – genetic gain- and loss-of function. **A)** Scheme of floxed MYC alleles, which can be deleted by Cre^ERT2^ recombinase upon treatment with 4-OHT (4-Hydroxytamoxifen). E1-E3: Exon1-Exon3; 4-OHT: 4-Hydroxytamoxifen. **B)** Protein expression of MYC and beta actin (loading control) in EtOH and 4-Hydroxytamoxifen PDAC cells 72h after treatment. Displayed are 3 independent biological replicates. **C)** Quantification of MYC protein expression, determined by western blot (n=3). **D)** Relative viability of PDAC cells, 72h after treatment with Bortezomib. Cells were pretreated with EtOH and 4-Hydroxytamoxifen for 24h. Viability was measured by MTT-test. P value of an unpaired t-test is depicted (n=3). **E)** Quantitative PCR of indicated targets 72h after treatment with 600nM 4-OHT. *Gapdh* served as housekeeping control (n=3). **F)** Viability test by CellTiterGlo of PSN1 and PPT-9091-MYC^ER^ cell lines. 2.000 cells were seeded and after 24h treated with 600nM 4-OHT (MYC^ER^ shuttles into nucleus) or EtOH (vehicle) and 7 increasing concentrations of Bortezomib for three days; highest conc.: 100nM. **J)** 6 days treatment with 600nM of 4-OHT, and simultaneous treatment with Bortezomib 24h after seeding of 1.000 cells/well similar to (H). **K)** Treatment for 3d with 600nM 4-OHT and subsequent 3d treatment with Bortezomib without 4-OHT treatment according to (I). For I-K the standard deviation (SD) was used for error bars and three independent biological replicates were conducted as technical triplicates.

## Discussion

Success of cancer therapeutics substantially differs due to a huge heterogeneity of human cancers, incomplete understanding how drugs mechanistically act, poorly described resistance mechanisms, or a lack of stratification for patients which benefit from the therapy. Here we performed a limited unbiased pharmacological screen and provide evidence that perturbants of the protein homeostasis are more effective in MYC hyperactive PDAC cells.

MYC is well known to serve the metabolic demands for biomass accumulation of dividing cells, including a prominent function towards protein synthesis through increasing ribosome biogenesis [49]. The relevance of MYC-induced protein synthesis for its function in cancer is well documented. Ribosomal protein haploinsufficiency impairs MYCs oncogenic activity in the *Eμ-Myc* lymphoma model [50]. High MYC activity increases the protein load beyond the protein folding capacity of cells and can therefore activate UPR in mammalians and Drosophila [51, 52]. The importance of MYC-induced UPR is underscored by the demonstration of a synthetic lethal interaction of MYC with components of the UPR, including PERK and XBP1 [51, 53, 54]. Across several PDAC data-sets and in mechanistic conditional MYC off/on models, we observed a connection of MYC activity to UPR signatures, arguing that PDACs with high MYC activity might be at the edge to die from proteotoxicity. Although such cancer cells can cope with the increased protein load via an adaptive ER-stress-induced survival pathway [37], they are less able to tolerate any further increased protein challenge, contributing to our observation of increased proteasome inhibitor sensitivity in at least some MYC-hyperactive PDACs. Such a scenario is supported by several layers of evidence. It was demonstrated that PDAC cells escaping dependency on KRAS activate the MYC network to increase protein synthesis, which activates adaptive ER-stress pathways [55]. Consistent with our data, such PDAC cells were found to be susceptible to perturbations of protein homeostasis induced by HSP90- or proteasome-inhibitors [55]. Moreover, we found that human PDAC lines sensitive to the Valosin-containing protein (VCP)/p97 inhibitor NMS-873, known to trigger a UPR [45, 46], enrich for MYC signatures. The strong connection of MYC to translation, the observed link to the UPR, the enrichment of MYC signatures in p97 inhibitor or HSP90 inhibitor sensitive PDAC cell lines, and the modulation of the proteasome inhibitor response in MYC genetic gain- and loss-of-function models [38], argues that MYC hyperactive PDACs are more sensitive towards perturbations of the protein homeostasis. Although these consideration need additional validations in context of PDAC, they are supported from clinical data in multiple myeloma, where MYC seem to be connected to a benefit of proteasome inhibitor based therapy [56, 57]. Some PDAC xenograft *in vivo* models respond to proteasome inhibitor treatment [58, 59], whereas others resist [60]. Also, for patient-derived xenotransplants (PdXs), proteasome inhibitor responding and non-responding models have been documented [61, 62]. Interestingly, Beglyarova et al. observed a proteasome inhibitor response in a MYC-amplified PdX with high protein expression of the oncogene, whereas the non-amplified PdX tested in the study resisted the therapeutic intervention [62]. Such observations clearly demonstrate the need to stratify for responsiveness towards perturbations of protein homeostasis and support our note that proteasome inhibitor sensitivity is a MYC-associated trait in PDAC. The lack of stratification might contribute to the negative outcome of a phase II PDAC study, where patients were treated with Bortezomib or with the combination containing gemcitabine and Bortezomib [63].

In the neuroblastoma line SHEP, which harbors a MYCN^ER^ transgene, an unbiased pharmacological screen with 938 FDA-approved drugs, recently demonstrated Bortezomib, carfilzomib, cabazitaxel, pralatrexate, gemcitabine, vincristine, docetaxel, paclitaxel, etoposide, and doxorubicin to be the top ten MYCN-associated pharmacological vulnerabilities [64]. The substantial overlap of these hits with our screen validates the used experimental approach and demonstrates specific vulnerabilities across the MYC family and cross different tumor entities. As in the PDAC context, where MYC directly activates the transcription of the pro-death BCL2 family member *NOXA* (*PMAIP1*) [38] upon Bortezomib treatment, NOXA contributes also in neuroblastoma models significantly to the Bortezomib-induced apoptosis [64].

The DoRothEA database [32] demonstrates that MYC has the highest number of transcription factor-drug interactions among all transcription factors analyzed [32]. Interestingly, only associations in which MYC is sensitizing for a drug were observed in this database [32]. Consistently, in context of PDAC, evidence that MYC increases the sensitivity towards proteasome inhibitors, BET inhibitors [27, 28], SUMOylation inhibitors [29], the ERCC3 inhibitor triptolide [62], or Cisplatin [65] was provided. However, it is important to note, that MYC was also associated with drug resistance in PDAC. Important work has demonstrated that MYC is involve in a ductal-neuroendocrine lineage switch, whereby the neuroendocrine lineage resist Gemcitabine [66]. Although paclitaxcel was demonstrated to trigger a mitotic vulnerability [67], recent work, which investigated paclitaxcel-resistant primary PDAC cultures implicates a MYC function in the resistant phenotype [68]. Interestingly, the anti-apoptotic BCL2 family member MCL1 is co-upregulated with MYC in paclitaxel-resistant PDAC cultures [68]. Anti-apoptotic BCL2 family members are described to be relevant modulators of the MYC associated mitotic vulnerability [67]. Whether BCL2 family proteins are important switches, determining MYC mediated sensitivity or resistance in the PDAC context awaits further detailed analysis.

As a mono- as well as in combination-therapies, Bortezomib demonstrates limited efficacy in solid cancers in the clinic [69–71]. Furthermore, a narrow therapeutic index and unfavorable pharmacokinetic features [69, 71], with impaired distribution to solid tumors, may limit the clinical development of Bortezomib in PDAC. However, our data provide evidence that perturbation of the protein homeostasis is an option to target MYC-active PDACs. Considering the development of next-generation proteasome inhibitors [69], the development of new bortezomib formulations [72], or options to target the ubiquitin-proteasome system at different levels [45, 46, 73], will allow to advance the concept in the future.

## Material and methods

### Analysis of publicly available expression data, drug sensitivity data and clinical data

RNA-expression data of pancreatic cancer cell lines, included in the CCLE dataset (19Q3), was downloaded from the depmap data portal (https://depmap.org/). Drug sensitivities of human PDAC cell lines from the PRISM repurposing primary screen (19Q3) [39], the GDSC2 screen (AUC) [40], and the CTD^2 (AUC) screen [41] were directly accessed and downloaded via the depmap data portal. Bortezomib sensitivity of the lines was divided into quartiles and the most sensitive quartile was investigated for pathway enrichment using gene set enrichment using the complete CCLE-PDAC dataset. For the analysis of drug-MYC interactions we accessed the DoRothEA (Discriminant Regulon Expression Analysis) database [32] (http://dorothea.opentargets.io/) and extracted significant (FDRq < 0.05) drug hits, which are sensitive in cells with an increased MYC expression. Drug hits were summarized in drug classes and compared with hits of our experimental drug screening in a Venn diagram.

PDAC transcriptome data sets of the cancer genome atlas (TCGA) were curated according to Peran et al. [34] (n=150) and mRNA expression data and clinical data were accessed via the GDC data portal (https://portal.gdc.cancer.gov/) [33]. The ICGC dataset was downloaded from the supplemental data of [7]. Acinar cell carcinomas and intraductal papillary mucinous neoplasms were excluded (n=81). TCGA and ICGC datasets were clustered using ClustVis [74] using Euclidean for distance and the Ward method. The datasets were clustered according to the genes of the HALLMARK-MYC-TARGET_V1, HALLMARK-MYC-TARGET_V2, and the direct MYC targets determined by Muhar et al. [31]. PDAC identified by all three signatures were considered as common MYC^high^ PDACs. For the TCGA dataset clustered by HALLMARK-MYC-TARGET_V1, a cluster with incomplete high expression of the target genes was recognized and included in the MYC^high^ group according to this gene set. Survival data was assigned to the commonly MYC^high^ PDAC subtype and displayed in a Kaplan-Meyer-Curve. For subtype association of the common MYC^high^ PDAC, the subtyping of Bailey at al. [7] was used and the Pancreatic progenitor subtype, ADEX subtype, and the Immunogenic subtype were combined and depicted as non-squamous. The TCGA dataset were subtyped according to the identifier published by Moffitt et al. [75]. RNA-seq data of untreated and 4-OHT treated IMIM-PC1^MYCER^ cells were described [29] and can be accessed via NCBI/GEO: GSE119423. Enrichment analysis of gene sets was performed using the gene set enrichment analysis (GSEA) tool with default parameters (weighted) depending on sample size version 4.0.3 with signatures of the Molecular Signatures Database v7.0 and the MYC target gene set from Muhar et al. [31]. The false discovery rate (FDR) q-values and normalized enrichment scores are depicted in the figures. The signature: ATF4 HUMAN TF ARCHS4 COEXPRESSION was downloaded via the EnrichR database [76]. In addition to the weighted GSEA we performed an unweighted analysis of gene sets using the web tool GeneTrail2 1.6 [30], multiple testing was corrected according to [77] and displayed as adjusted p-value.

### Cell lines, CRISPR/Cas9 mediated knockout

Cell lines were cultured in high glucose DMEM medium (Life Technologies, Darmstadt, Germany) or RPMI (Life Technologies, Darmstadt, Germany) supplemented with 10% (w/v) fetal calf serum (FCS) (Merck Millipore, Berlin, Germany) and 1% (w/v) penicillin/streptomycin (Life technologies, Darmstadt, Germany). All murine pancreatic cancer cell lines were established from Kras^G12D^-driven mouse models of pancreatic cancer and cultivated as described [78]. Identity of the murine pancreatic cancer cell lines was verified by genotyping PCR. All human cell lines (Panc1, DanG, PaTu8988S, PSN1, PaTu8988T, MiaPaCa-2, IMIM-PC1) were authenticated by Multiplexion (Multiplexion GmbH, Heidelberg, Germany). To screen for Mycoplasma contamination all cell lines are tested by PCR as described [79]. The dual recombinase system [47] was used to generate a murine PDAC cell line allowing to delete floxed *Myc* alleles [48] by a tamoxifen activatable Cre (CRE^ERT2^). Alleles and genotyping for this murine PDAC cell line were recently described [80] and the PPT-MW1955 line corresponds to the following genotype: *Pdx1-Flp;FSF-Kras^G12D/+^;FSF-R26^CAG-CreERT2/+^;Myc^lox/lox^.* The murine cell line PPT-9091 transduced with the pBabepuro-myc-ER construct, which was a gift from Wafik El-Deiry (Addgene plasmid # 19128 ; http://n2t.net/addgene:19128 ; RRID:Addgene_19128) as described [29]. IMIM-PC1MYCER cells were described recently [29].

To generate the CRISPR/Cas9 mediated NOXA knockout the protein coding region of NOXAs exon two was targeted by two sgRNAs (sg#1: T C G A G T G T G C T A C T C A A C T C; sg#2: T G T A A T T G A G A G G A A T G T G A), which were cloned into the pKLV-U6gRNA(BbsI)-PGKpuro2ABFP vector which was a gift from Kosuke Yusa (Addgene plasmid # 50946; http://n2t.net/addgene:50946; RRID: Addgene_50946). MiaPaCa2 cells were co-transfected with a Cas9 expressing px330 vector and the two guides or the pKLV backbone only. Positive transfected MiaPaCa-2 cells were grown under puromycin treatment (1μg/ml) for two weeks. Subsequently single clones were generated, isolated and screened via PCR for knockout clones. The primer-set: C A C T A G T G T G G G C G T A T T A G G (FW) + G A T G T A T T C C A T C T T C C G T T T C C (RV1) reveals a product of 157 bp for knockout cells and 342 bp for wild type cells (data not shown). To further test if both alleles are deleted the primer set: FW + G T T C A G T T T G T C T C C A A A T C T C C (RV2) was used; here a product at 137bp is amplified if the cells harbor a NOXA allele and no product if the cells harbor a knockout for NOXA (Fig. 4C).

### Cell lysis and Western blot

To prepare whole-cell extracts RIPA buffer (50 mM Tris, 150 mM NaCl, 1% Nonidet P-40, 0.5% sodium deoxycholate, 0.1% SDS, pH 8.0) supplemented with protease and phosphatase inhibitors (Protease inhibitor cocktail complete EDTA free, Roche Diagnostics, Mannheim, Germany and Phosphatase-Inhibitor-Mix I, Serva, Heidelberg, Germany) was used. Whole-cell extracts were normalized for protein and heated at 95°C for 5 min in protein loading buffer (45.6 mM Tris-HCl pH 6.8, 2% SDS, 10% glycerol, 1% β-mercaptoethanol, 0.01% bromophenol blue) and loaded onto 10-12% SDS-polyacrylamide gels and proteins were transferred to nitrocellulose membranes (Merck-Millipore, Berlin, Germany). Afterwards, membranes were blocked in blocking buffer (5% skim milk, 0.1% Tween in PBS) and incubated with β-Actin (#A5316, Sigma-Aldrich), MYC (#9402, Cell Signaling) and cleaved PARP (552596, BD Pharmingen) primary antibodies. After overnight incubation (4°C) with primary antibodies, membranes were incubated with DyLight^TM^ 680 (#5366S, Cell Signaling) or 800 (#5151S, Cell Signaling) conjugated secondary antibodies (1:10000 dilution). Western blots were visualized by the Odyssey Infrared Imaging System (Licor, Bad Homburg, Germany) and quantified using the Image Studio Lite Software V 5.2.5 (Licor, Bad Homburg, Germany). Cleaved PARP and MYC expression values were normalized on β-Actin expression and final expression values were calculated out of 3 biological replicates.

### Quantitative real time PCR

To isolate RNA from cell lines we followed the manufacturer's instructions of the Maxwell 16 LEV simply RNA Kit (# AS1280) (Promega, Walldorf, Germany). Quantification of mRNA was performed using the BRYT Green® Dye (GoTaq® qPCR, #A600A, Promega) in a real-time PCR analysis system (StepOnePlus, Real-Time PCR System; Applied Biosystems Inc., Carlsbad, CA, USA). Primers used (5’-3’): *Myc*: T T C C T T T G G G C G T T G G A A A C (FW) / G C T G T A C G G A G T C G T A G T C G (RV), *Odc1*: A C A T C C A A A G G C A A A G T T G G (FW) / A G C C T G C T G G T T T T G A G T G T (RV), *Cad*: C T G C C C G G A T T G A T T G A T G T C (FW) / G G T A T T A G G C A T A G C A C A A A C C A (RV) *Gapdh*: G G G T T C C T A T A A A T A C G G A C T G C (FW) / T A C G G C C A A A T C C G T T C A C A (RV). Data analysis was carried out with StepOne software 2.3 Life Technologies/Applied Biosystems/ThermoFisher, Darmstadt, Germany) by the ΔΔCt method (as housekeeping gene *Gapdh* was used) as described [29].

### Compounds

The anti-cancer compound library with n=129 drugs was obtained as a plated compound set from the NCI/DTP Open Chemicals Repository (NCI/DTP, MD, USA); the full list of compounds is shown in supplemental table 3. Bortezomib was purchased from LC-Laboratories (Woburn, MA, USA), Marizomib was purchased from Cayman Chemicals (Ann Arbor, MI, USA) and 4-Hydroxytamoxifen (4-OHT) was purchased from Sigma (Sigma, Munich, Germany).

### Drug screening experiment

For the drug screen we adapted a recent screening approach [81]. In an attempt to select for drugs highly active in PDAC, we screened PaTu-8988S, Panc1, DanG, and PSN1 cells with a single dose of 600 nM of each drug. Screening was conducted in a 96 well format. 24 hours after the seeding (3.000 cells per well), cells were treated with the drugs of the anti-cancer compound library for additional 72 hours. Afterwards viability was measured with MTT assays as a read-out for the responsivity. The screen was performed as biological triplicates conducted as technical triplicates. The mean response in the MYC^high^ models was divided by the mean response in the MYC^low^ models. Drugs were ranked according to the ratio and a ratio >2 was defined as a hit.

### Viability assay and Clonogenic assay

38 recently characterized [42] murine PDAC cell lines driven by *Kras^G12D/+^* were termed as PDAC KC cell lines. Cell lines were seeded in a 96 well format. 24 hours after the seeding (1.500 cells per well), cells were treated with the respective drugs for additional 72 hours. Afterwards viability was measured with MTT assays as a read-out for the responsivity. 3-(4,5-Dimethylthiazol-2-yl)-2,5-diphenyltetrazolium bromide (Sigma, Munich, Germany) was used in a dilution of 5 mg/ml. 10 μL of this MTT solution was added per well and incubated for four hours at 37°C. Subsequently the medium was removed and the formazan crystals dissolved in 200 μL DMSO:EtOH (v/v) and incubated for 10 minutes on a horizontal shaker. Absorption was measured at 595 nm on a Thermo/LabSystem Multiskan RC Microplate Reader (Artisan Technology Group, Champaign, IL, USA).

In addition to MTT assay, cellular viability was measured by CellTiter-Glo ATP Viability Assay. Briefly, 25 μl CellTiter-Glo® reagent purchased from Promega (Fitchburg, WI, USA) was added to each well of a 96-well plate 72 hours after drug treatment. After 10 minutes of gentle shaking and 20 minutes of incubation at room temperature, luminescence was measured on a FLUOstar OPTIMA microplate reader (BMG Labtech Ortenberg, Germany). The growth inhibitory 50% (GI_50_) concentration was calculated with GraphPad Prism 6 (GraphPad Software, CA, USA) using a non-linear regression model. For the clonogenic assay 2.000 MiaPaCa-2 cells (wildtype or NOXA knockout) were seeded in 12-well plates. After 24 hours cells were treated once with the indicated doses of Bortezomib followed by culturing for 14 days in DMEM medium (Life Technologies, Darmstadt, Germany) supplemented with 10% (w/v) fetal calf serum (FCS) (Merck Millipore, Berlin, Germany) and 1% (w/v) penicillin/streptomycin (Life technologies, Darmstadt, Germany). Afterwards the medium was carefully removed, and cells were washed three times with PBS. The colonies were stained with 0.2% Crystal Violet solution (Sigma by Life Technologies TM, Darmstadt, Germany) for 20 minutes on a shaker at room temperature. To remove background staining, the wells were washed 3 times with tap water, dried and subsequently visualized using a flatbed scanner.

### Statistical methods

All experiments were conducted in biological triplicated unless otherwise stated in the figure legends. ANOVA or two-sided Student’s t-test was used to investigate statistical significance, as indicated. p-values were calculated with GraphPad Prism6/8 (GraphPad Software, CA, USA) and corrected according to Bonferroni for multiple testing unless otherwise indicated. p-values are indicated or * in the figures denotes p < 0.05. Fisher’s exact test, was used to assess the association between PDAC subtypes and the expression of MYC target genes.

## Abbrevations

4-OHT: 4-hydoxytamoxifen
ATF4: activating transcription factor 4
BET: bromodomain and extra terminal motif
CNV: copy number variation
CTD2: cancer target discovery and development network
depmap: dependency map
DoRothEA: discriminant regulon expression analysis
GSEA: gene set enrichment analysis
ICGC: international cancer gene consortium
MYC: myelocytomatosis oncogene
PDAC: pancreatic ductal adenocarcinoma
PERK: proteinkinase RNA-activated-like ER kinase
SUMO: small-ubiquitin-like modifier
TCGA: the cancer genome atlas
UPR: unfolded protein response
UPS: ubiquitin proteasome system

## Acknowledgment

We thank NCI/DTP Open Chemicals Repository for providing the screening library. The results published here are fully or partially based upon data generated by the Cancer Target Discovery and Development (CTD^2^) Network (https://ocg.cancer.gov/programs/ctd2/data-portal) established by the National Cancer Institute’s Office of Cancer Genomics. Part of the results shown here are based upon data generated by the TCGA Research Network: https://cancergenome.nih.gov. We thank all colleagues providing plasmids via the Addgene platform. We thank Aylin Aydemir for excellent technical support.

## Supplementary Information

**Table S1** *Doubling time of PDAC cell lines used for the drug screening experiment.* Panc1, PaTu8988S, DanG and PSN1 cells were seeded in 96 well plates at a density of 3000 cells/well. After 24, 48h, 72h and 96h viable cells were measured by MTT-test to determine doubling time of the cell lines.

**Table S2** *Complete list of the response to compounds used in the drug screen.* PSN1, Panc1, PaTu8988S and DanG cells were treated with n=129 compounds in biological and technical triplicates. Drug response as well as mean of MYC high and mean of MYC low are displayed.

**Table S3** *Pathways associated with MYC in common MYC^high^ PDAC.* GSEA of pathways enriched in common MYC^high^ PDAC of the ICGC and the TCGA dataset.

**SFigure 1** *Survival and Subtypes of common MYC^high^ PDACs.* **A)** Survival data of common MYC^high^ PDACs of the ICGC dataset are displayed in a Kaplan-Meyer-Curve. P value of a log-rank test is depicted. **B)** Percentage of squamous subtype of the common MYC^high^ PDACs compared to the others. Fisher Exact test: p<0.0001. **C)** Venn diagram of PDAC identified as MYC^high^ by clustering of the genes of the depicted signatures in the TCGA dataset. 16 PDACs were identified as common MYC^high^ PDACs. **D)** Survival data of common MYC^high^ PDACs of the TCGA dataset are displayed in a Kaplan-Meyer-Curve. **E)** Percentage of basal-like subtype of the common MYC^high^ PDACs compared to the others. Fisher Exact test: p<0.05. F) GSEA of IMIM-PC1^MYCER^ cells treated with 4-OHT to activate MYC. Depicted are HALLMARK and KEGG signature corresponding to the tissue-based analysis corresponding to Fig. 3D. The NES and the FDR q values are depicted.

**SFigure 1** *Association of MYC with perturbants of the protein homeostasis.* Sensitivities of human PDAC cell lines from the PRISM repurposing primary screen (19Q3) [39] of the depicted drug classes were divided into quartiles and lines for the most sensitive quartile were compared to the remaining cell lines of the complete CCLE PDAC dataset with a gene set enrichment analysis using the GeneTrail2 1.6 web service. The enrichment score was color coded. ** adjust. p-value<0.01; **** adjust. p-value<0.0001.

