## Supplemental Files for "Targeting the ubiquitin-proteasome system in a pancreatic cancer subtype with hyperactive MYC"

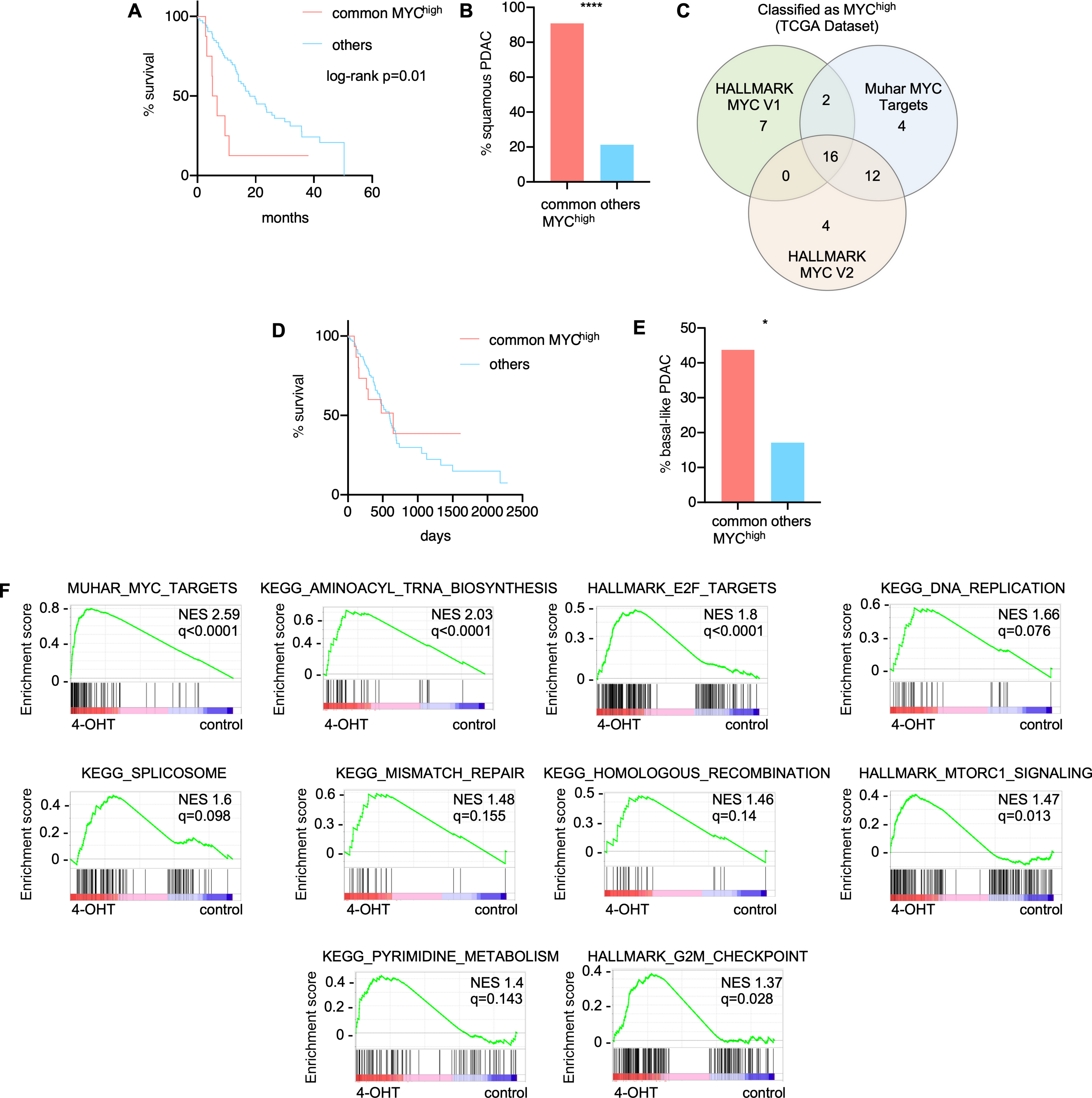
A
common MYC high
others
log-rank p=0.01
% survival
months
B
****
% squamous PDAC
common MYC high
others
C
Classified as MYC high (TCGA Dataset)
HALLMARK MYC V1
Muhar MYC Targets
HALLMARK MYC V2
D
common MYC high
others
% survival
days
E
*
% basal-like PDAC
common MYC high
others
F
MUHAR_MYC_TARGETS
NES 2.59 q<0.0001
KEGG_AMINOACYL_TRNA_BIOSYNTHESIS
NES 2.03 q<0.0001
HALLMARK_E2F_TARGETS
NES 1.8 q<0.0001
KEGG_DNA_REPLICATION
NES 1.66 q=0.076
KEGG_SPLICOSOME
NES 1.6 q=0.098
KEGG_MISMATCH_REPAIR
NES 1.48 q=0.155
KEGG_HOMOLOGOUS_RECOMBINATION
NES 1.46 q=0.14
HALLMARK_MTORC1_SIGNALING
NES 1.47 q=0.013
KEGG_PYRIMIDINE_METABOLISM
NES 1.4 q=0.143
HALLMARK_G2M_CHECKPOINT
NES 1.37 q=0.028
Enrichment score
4-OHT
control

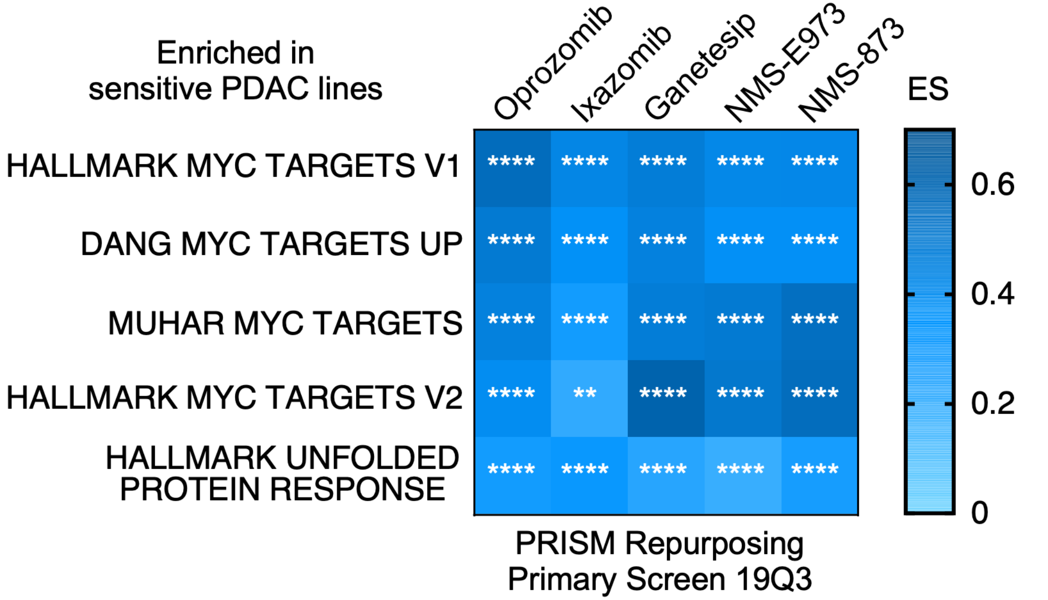
Enriched in
sensitive PDAC lines
Oprozomib
Ixazomib
Ganetesip
NMS-E973
NMS-873
ES
HALLMARK MYC TARGETS V1
DANG MYC TARGETS UP
MUHAR MYC TARGETS
HALLMARK MYC TARGETS V2
HALLMARK UNFOLDED
PROTEIN RESPONSE
****
**
0.6
0.4
0.2
0
PRISM Repurposing
Primary Screen 19Q3

| cell line ID | doubling time (h) |
|---|---|
| Panc1 | 44.78 |
| Patu8988S | 48.86 |
| DanG | 45.62 |
| PSN1 | 27.31 |

| | Panc1 | Patu8988S | DanG | PSN1 | |
|---|---|---|---|---|---|
| | Myc low | Myc low | Myc high | Myc high | Mean MYC high / Mean MYC low |
| Panobinostat | 0.5927368 | 0.4316097 | 0.1283566 | 0.1012438 | 4.461432123 |
| Cladribine | 1.0362948 | 0.8812915 | 0.3784469 | 0.1964837 | 3.335335254 |
| Idarubicin hydrochloride | 0.3853088 | 0.3580649 | 0.1316619 | 0.106136 | 3.126073667 |
| Clofarabine | 0.6907533 | 0.9303737 | 0.3631497 | 0.1735106 | 3.020768958 |
| Topotecan hydrochloride | 0.6338552 | 0.7889655 | 0.3766401 | 0.146818 | 2.718118017 |
| Gemcitabine hydrochloride | 0.6285325 | 0.7472625 | 0.3121124 | 0.2189043 | 2.590869276 |
| Floxuridine | 0.7672699 | 0.9213013 | 0.489905 | 0.3033217 | 2.128737275 |
| Doxorubicin hydrochloride | 0.5381085 | 0.4160912 | 0.309046 | 0.1406154 | 2.122040469 |
| Bortezomib | 0.1740942 | 0.3174964 | 0.132343 | 0.1032272 | 2.086811521 |
| Dactinomycin | 0.2778748 | 0.4070966 | 0.2398357 | 0.0918611 | 2.065052824 |
| Pralatrexate | 0.5334845 | 0.8566843 | 0.3301672 | 0.3439406 | 2.062235104 |
| Daunorubicin hydrochloride | 0.3201861 | 0.6879631 | 0.4281178 | 0.084397 | 1.967063898 |
| Dasatinib | 1.0649763 | 0.8051341 | 0.3248287 | 0.6516466 | 1.915164076 |
| Methotrexate | 0.543753 | 0.8269723 | 0.3974606 | 0.3274105 | 1.890991486 |
| Romidepsin | 0.6142233 | 0.6397273 | 0.4472008 | 0.2266042 | 1.860999368 |
| Pemetrexed, Disodium salt, Heptahydrate | 1.1160514 | 0.8835727 | 0.7531286 | 0.3413614 | 1.826991624 |
| Ixabepilone | 0.6076967 | 0.4359349 | 0.1981229 | 0.3851835 | 1.789165199 |
| Epirubicin hydrochloride | 0.5976475 | 0.600278 | 0.4432164 | 0.2435132 | 1.744391794 |
| Paclitaxel | 0.5937123 | 0.4850653 | 0.2299463 | 0.3899297 | 1.740312281 |
| Mitomycin | 0.8186603 | 0.7457391 | 0.5472753 | 0.3686277 | 1.708040398 |
| Trametinib | 1.0522746 | 0.3858011 | 0.5152489 | 0.3443984 | 1.672867182 |
| Belinostat | 1.2235609 | 0.9651008 | 0.6105508 | 0.742512 | 1.617561069 |
| Plicamycin | 0.2406453 | 0.5661695 | 0.3741679 | 0.1455876 | 1.552296875 |
| Docetaxel | 0.6006985 | 0.358746 | 0.203751 | 0.4255217 | 1.524688004 |
| Cytarabine hydrochloride | 0.7906227 | 0.7922291 | 0.4905566 | 0.5891065 | 1.466060789 |
| Valrubicin | 0.658399 | 0.6932424 | 0.6135694 | 0.3178223 | 1.4512062 |
| Vincristine sulfate | 0.4842991 | 0.4904168 | 0.3563123 | 0.3683089 | 1.34513844 |
| Cobimetinib | 0.8057013 | 0.4227304 | 0.5749653 | 0.3467257 | 1.332801963 |
| Cabazitaxel | 0.5594477 | 0.4260106 | 0.3483661 | 0.3990782 | 1.318437226 |
| Vinblastine sulfate | 0.4661904 | 0.6971546 | 0.5479773 | 0.3792634 | 1.254630907 |
| Teniposide | 0.7099909 | 0.6541106 | 0.5130224 | 0.5785205 | 1.249700352 |
| Vorinostat | 1.1984255 | 1.0365938 | 0.8138551 | 0.985923 | 1.241830561 |
| Mitoxantrone | 0.4111703 | 0.360539 | 0.4180332 | 0.2049134 | 1.23880501 |
| Carfilzomib | 0.123085 | 0.2872714 | 0.1195335 | 0.2173406 | 1.218130104 |
| Pazopanib hydrochloride | 1.1507001 | 1.1699473 | 0.9319465 | 0.9738086 | 1.217704961 |
| Tamoxifen citrate | 1.2258223 | 0.8767898 | 0.8112547 | 0.9373592 | 1.202445098 |
| Vemurafenib | 1.2983508 | 1.0166108 | 0.9096793 | 1.0162834 | 1.201976341 |
| Bleomycin sulfate | 0.9200904 | 0.8969911 | 0.7066424 | 0.8128505 | 1.195847316 |
| Axitinib | 1.0551648 | 1.0017045 | 0.816527 | 0.9102405 | 1.191167527 |
| Thioguanine | 1.0981832 | 0.8629344 | 1.0158377 | 0.6383698 | 1.185533008 |
| Vinorelbine tartrate | 0.5201178 | 0.5633463 | 0.497125 | 0.4253102 | 1.174569397 |
| Sorafenib | 1.162171 | 1.2830505 | 1.0178113 | 1.0745961 | 1.168616375 |
| Lenvatinib | 1.149886 | 1.0538571 | 0.8996172 | 0.9908693 | 1.165701528 |
| Ponatinib | 0.9676809 | 0.8053608 | 0.730816 | 0.8036482 | 1.155479361 |
| Trifluridine | 1.1512183 | 1.0878322 | 0.9053789 | 1.0408002 | 1.150485363 |
| Pomalidomide | 1.0906291 | 1.0283482 | 0.9547646 | 0.9212327 | 1.129520454 |
| Celecoxib | 1.1502735 | 1.1248195 | 0.9611498 | 1.0822925 | 1.113362995 |
| Idelalisib | 1.1145138 | 1.0326372 | 0.9856933 | 0.9471336 | 1.110886312 |
| Anastrozole | 1.0940645 | 1.0583861 | 0.9622566 | 0.9757917 | 1.110627962 |
| Megestrol acetate | 1.1106453 | 1.1060051 | 0.9991441 | 1.0044367 | 1.106344418 |
| Mechlorethamine hydrochloride | 1.1069868 | 0.9741389 | 1.0350162 | 0.8476075 | 1.105438979 |
| Raloxifene | 1.0973425 | 1.055331 | 1.0202812 | 0.9277671 | 1.1050411 |
| Chlorambucil | 1.2870682 | 1.0877594 | 1.0875412 | 1.0623749 | 1.104614114 |
| Omacetaxine mepesuccinate | 0.1392018 | 0.5614243 | 0.4482203 | 0.1877929 | 1.101590466 |
| Carboplatin | 1.1176001 | 1.0182414 | 0.9936223 | 0.9558329 | 1.095609456 |
| Regorafenib | 1.1063028 | 0.9268283 | 0.9606536 | 0.8963906 | 1.094821059 |
| Decitabine | 0.9667371 | 0.9534292 | 0.622724 | 1.1400811 | 1.089267518 |
| Afatinib | 1.0705093 | 0.892516 | 0.8782874 | 0.9339288 | 1.083218081 |
| Hydroxyurea | 1.1290124 | 0.9838368 | 0.9961748 | 0.9549181 | 1.082905492 |
| Amifostine | 1.0944602 | 1.0792695 | 0.9797668 | 1.0289677 | 1.082138826 |
| Fluorouracil | 1.0876493 | 1.0280319 | 0.9256822 | 1.0320756 | 1.08066543 |
| Mitotane | 1.1561005 | 1.0283327 | 1.0450807 | 0.9784554 | 1.079512906 |
| Crizotinib | 1.009358 | 0.8524901 | 0.7347524 | 0.9914502 | 1.078580271 |
| Carmustine | 1.1694594 | 1.0111272 | 1.0154788 | 1.0152511 | 1.073794532 |

| | | | | | |
|---|---|---|---|---|---|
| Lenalidomide | 1.1758563 | 1.0865372 | 0.9625862 | 1.1516524 | 1.070074789 |
| Arsenic trioxide | 1.1543637 | 1.0026565 | 1.0099186 | 1.0098961 | 1.067929769 |
| Dabrafenib mesylate | 1.0476995 | 0.7038174 | 0.6271765 | 1.0143573 | 1.067000238 |
| Streptozocin | 1.0629211 | 1.1350681 | 1.0200487 | 1.0443164 | 1.064728879 |
| Estramustine phosphate sodium | 1.1174815 | 0.9156263 | 0.9230253 | 0.9915053 | 1.061935398 |
| Aminolevulinic acid hydrochloride | 1.2735396 | 1.0125003 | 1.0472284 | 1.1059952 | 1.061682585 |
| Vismodegib | 1.1140285 | 1.0051374 | 0.978548 | 1.0176179 | 1.061618157 |
| Methoxsalen | 1.1077994 | 1.0322965 | 1.0017812 | 1.017235 | 1.059969685 |
| Uridine triacetate | 1.1105191 | 0.934566 | 0.9040666 | 1.026801 | 1.05915346 |
| Ifosfamide | 1.0673416 | 1.0517827 | 0.9984629 | 1.0063229 | 1.057032755 |
| Pentostatin | 1.1168985 | 1.1328357 | 1.0000841 | 1.1287894 | 1.056772122 |
| Etoposide | 0.8257375 | 0.9160592 | 0.8008493 | 0.8583683 | 1.049769979 |
| Plerixafor | 1.0156081 | 1.0299569 | 0.9693547 | 0.9840821 | 1.047162072 |
| Mercaptopurine | 1.1533893 | 1.064621 | 1.0421167 | 1.0762546 | 1.047035651 |
| Temozolomide | 1.100926 | 0.9987936 | 1.0170777 | 0.9943394 | 1.043900587 |
| Melphalan hydrochloride | 1.1167922 | 0.8871951 | 0.9884728 | 0.9317933 | 1.043598772 |
| Gefitinib | 1.0671326 | 0.9157831 | 0.9355389 | 0.9649645 | 1.043363337 |
| Busulfan | 1.1515438 | 0.9183881 | 1.019652 | 0.9703823 | 1.040148863 |
| Exemestane | 1.0891907 | 1.0271264 | 1.0561781 | 0.9864502 | 1.036075461 |
| Altretamine | 1.2215769 | 0.9396757 | 1.0638617 | 1.0242761 | 1.035014345 |
| Triethylenemelamine | 0.8480403 | 0.8116295 | 0.6936217 | 0.9105794 | 1.034577136 |
| Ibrutinib | 1.0167125 | 0.8967658 | 0.9005192 | 0.9503861 | 1.033806651 |
| Temsirolimus | 0.7991547 | 0.7438944 | 0.5569734 | 0.9390997 | 1.031399575 |
| Pipobroman | 1.0421908 | 1.0351135 | 0.9131288 | 1.1086645 | 1.027456309 |
| Cyclophosphamide | 1.0963373 | 1.0779427 | 1.0257545 | 1.0921417 | 1.026622568 |
| Nelarabine | 1.0364137 | 1.0449675 | 0.9784018 | 1.0608611 | 1.020653647 |
| Dacarbazine | 1.0773743 | 0.9799451 | 0.9998566 | 1.0329581 | 1.012054519 |
| Lapatinib | 1.2118227 | 1.0117246 | 1.0453586 | 1.1529822 | 1.011466159 |
| Olaparib | 1.0333574 | 0.9618045 | 0.9874746 | 0.9953946 | 1.006199381 |
| Azacitidine | 1.1057188 | 0.8669931 | 0.8983152 | 1.0624277 | 1.006104312 |
| Sunitinib | 1.125673 | 0.9359795 | 1.0082527 | 1.0437547 | 1.004700285 |
| Procarbazine hydrochloride | 1.0467224 | 0.9620981 | 0.9780714 | 1.0236254 | 1.003558771 |
| Cisplatin | 0.9783246 | 0.9864485 | 0.9717884 | 0.9881982 | 1.002442154 |
| Thalidomide | 1.1016887 | 0.8760751 | 0.9574126 | 1.0155748 | 1.002420886 |
| Capecitabine | 1.1033576 | 0.9785461 | 0.991254 | 1.087819 | 1.001361478 |
| Fludarabine phosphate | 1.0735069 | 0.9572477 | 1.0175318 | 1.0113273 | 1.000934311 |
| Letrozole | 0.964071 | 1.0074587 | 0.9956723 | 0.9810257 | 0.997385394 |
| Ixazomib citrate | 0.245811 | 0.5349317 | 0.3633141 | 0.4205316 | 0.996041413 |
| Erismodegib | 1.101577 | 0.8690601 | 0.9034578 | 1.0784045 | 0.99433599 |
| Erlotinib hydrochloride | 1.0831351 | 0.9835468 | 1.0028445 | 1.0771681 | 0.993591089 |
| Imatinib | 1.0910148 | 0.9407156 | 1.0124131 | 1.0327366 | 0.993438465 |
| Irinotecan hydrochloride | 1.0565756 | 0.8007072 | 0.9332201 | 0.9427394 | 0.990044197 |
| Tretinoin | 1.0659355 | 0.9976059 | 1.0807142 | 1.0058703 | 0.988956574 |
| Dexrazoxane | 1.0527764 | 0.9693531 | 0.9693255 | 1.0795052 | 0.986967628 |
| Uracil mustard | 1.001703 | 1.0043368 | 1.0454801 | 0.9906094 | 0.98524151 |
| Allopurinol | 1.1122121 | 1.0559206 | 1.0270413 | 1.1792553 | 0.982702291 |
| Vandetanib | 1.0904952 | 0.819051 | 0.9319007 | 1.0114376 | 0.98261128 |
| Bendamustine hydrochloride | 1.1159631 | 0.9338939 | 1.078548 | 1.0149397 | 0.979158848 |
| Sirolimus | 0.787637 | 0.736924 | 0.624014 | 0.9441733 | 0.972180429 |
| Enzalutamide | 0.9712496 | 0.9460363 | 1.008043 | 0.9676428 | 0.970440678 |
| Nilotinib | 1.1224254 | 1.146312 | 1.2644867 | 1.085551 | 0.965404678 |
| Thiotepa | 0.9899792 | 0.9029347 | 0.9891108 | 0.9768673 | 0.962835683 |
| Everolimus | 0.8114723 | 0.776071 | 0.6802286 | 0.9714906 | 0.961145951 |
| Lomustine | 0.9521525 | 0.9542411 | 1.0056073 | 0.9788209 | 0.960676527 |
| Oxaliplatin | 1.0346028 | 0.8738007 | 1.006481 | 0.9801285 | 0.960633421 |
| Cabozantinib | 1.0326035 | 0.8079131 | 0.8756074 | 1.0459597 | 0.95782062 |
| Palbociclib | 0.8332184 | 0.7235895 | 0.7367916 | 0.8895787 | 0.957228421 |
| Fulvestrant | 1.0301407 | 0.8667615 | 1.0025988 | 0.9857989 | 0.953985335 |
| Zoledronic acid | 1.1484752 | 0.9349664 | 1.1061776 | 1.0915123 | 0.94801437 |
| Osimertinib | 1.0508892 | 0.7746164 | 0.9351702 | 0.997295 | 0.94465117 |
| Alectinib | 0.9845287 | 0.765287 | 0.8784731 | 0.9897483 | 0.936621134 |
| Ceritinib | 0.9920572 | 0.6634162 | 0.9385272 | 0.8786845 | 0.910996414 |
| Bosutinib | 1.1090965 | 0.7804824 | 1.0985411 | 0.9776483 | 0.910118767 |
| Imiquimod | 1.0991822 | 0.8491946 | 1.2026013 | 0.9917523 | 0.887904663 |
| Abiraterone | 1.0443955 | 0.846116 | 1.1087497 | 1.1029108 | 0.854792766 |

| Signature | ICGC | NES | NOM p-val | FDR q-val | T |
|---|---|---|---|---|---|
| GO_2_IRON_2_SULFUR_CLUSTER_BINDING | | 1.5546366 | 0.04752475 | 0.12653017 | |
| GO_4_IRON_4_SULFUR_CLUSTER_BINDING | | 1.8781075 | 0.00431035 | 0.02116238 | |
| GO_90S_PRERIBOSOME | | 1.8662984 | 0.00609756 | 0.02314922 | |
| GO_AEROBIC_ELECTRON_TRANSPORT_CHAIN | | 1.6586405 | 0.02335456 | 0.07452937 | |
| GO_AEROBIC_RESPIRATION | | 1.9587013 | 0.01890756 | 0.01483259 | |
| GO_AMINO_ACID_ACTIVATION | | 2.0083132 | 0.00424629 | 0.01287796 | |
| GO_ANAPHASE_PROMOTING_COMPLEX | | 1.7914375 | 0.00212314 | 0.03398622 | |
| GO_ANAPHASE_PROMOTING_COMPLEX_DEPENDENT_CATABOLIC_PROCESS | | 1.8996791 | 0 | 0.01870802 | |
| GO_ANTIGEN_PROCESSING_AND_PRESENTATION_OF_EXOGENOUS_PEPTIDE_ANTIGEN_VIA_MHC_CLASS_I | | 1.5702072 | 0.09255533 | 0.11812779 | |
| GO_ANTIGEN_PROCESSING_AND_PRESENTATION_OF_PEPTIDE_ANTIGEN_VIA_MHC_CLASS_I | | 1.5075994 | 0.12525253 | 0.1587431 | |
| GO_APOPTOTIC_MITOCHONDRIAL_CHANGES | | 1.6247365 | 0.01310044 | 0.08969446 | |
| GO_ATP_SYNTHESIS_COUPLED_ELECTRON_TRANSPORT | | 1.9090444 | 0.00847458 | 0.01781047 | |
| GO_BASE_EXCISION_REPAIR | | 1.7991501 | 0.00662252 | 0.03281886 | |
| GO_CAJAL_BODY | | 1.7672838 | 0.00672646 | 0.03885208 | |
| GO_CATALYTIC_ACTIVITY_ACTING_ON_A_RRNA | | 1.7463319 | 0.01276596 | 0.04473388 | |
| GO_CATALYTIC_ACTIVITY_ACTING_ON_DNA | | 1.9750601 | 0.00428266 | 0.01349506 | |
| GO_CATALYTIC_ACTIVITY_ACTING_ON_RNA | | 2.0996804 | 0.00422833 | 0.00722887 | |
| GO_CATALYTIC_STEP_2_SPLICEOSOME | | 2.189515 | 0.00215517 | 0.00373597 | |
| GO_CELL_CYCLE_DNA_REPLICATION | | 1.8440794 | 0.00858369 | 0.02634763 | |
| GO_CELL_CYCLE_G2_M_PHASE_TRANSITION | | 1.9311495 | 0.00662252 | 0.01691992 | |
| GO_CELL_REDOX_HOMEOSTASIS | | 1.4075692 | 0.12103175 | 0.24583554 | |
| GO_CELLULAR_COMPONENT_DISASSEMBLY | | 1.6931647 | 0.00617284 | 0.06115709 | |
| GO_CELLULAR_METABOLIC_COMPOUND_SALVAGE | | 1.5438282 | 0.02964427 | 0.1331458 | |
| GO_CELLULAR_PROTEIN_COMPLEX_DISASSEMBLY | | 1.9894688 | 0.00213675 | 0.01319396 | |
| GO_CELLULAR_RESPIRATION | | 1.9116265 | 0.01301519 | 0.01747709 | |
| GO_CELLULAR_RESPONSE_TO_OXYGEN_LEVELS | | 1.5694808 | 0.04661017 | 0.11812197 | |
| GO_CHAPERONE_BINDING | | 1.4569552 | 0.06029106 | 0.20116156 | |
| GO_CHAPERONE_COMPLEX | | 1.8484074 | 0.0059761 | 0.02586903 | |
| GO_CHAPERONE_MEDIATED_PROTEIN_FOLDING | | 1.646454 | 0.0349076 | 0.07907362 | |
| GO_CHROMOSOME_SEPARATION | | 1.8884473 | 0.00643777 | 0.01995567 | |
| GO_CIS_TRANS_ISOMERASE_ACTIVITY | | 1.6616777 | 0.02531646 | 0.07315822 | |
| GO_CLEAVAGE_INVOLVED_IN_RRNA_PROCESSING | | 1.927207 | 0 | 0.01706572 | |
| GO_CONDENSED_CHROMOSOME_CENTROMERIC_REGION | | 1.908657 | 0.00212766 | 0.0177083 | |
| GO_COTRANSLATIONAL_PROTEIN_TARGETING_TO_MEMBRANE | | 1.8328004 | 0.01372549 | 0.02758209 | |
| GO_CYTOCHROME_COMPLEX | | 1.6537461 | 0.04408818 | 0.07639154 | |
| GO_CYTOCHROME_COMPLEX_ASSEMBLY | | 1.8497947 | 0.00214133 | 0.02585637 | |
| GO_CYTOPLASMIC_TRANSLATION | | 2.0662992 | 0.0060241 | 0.00829011 | |
| GO_CYTOPLASMIC_TRANSLATIONAL_INITIATION | | 1.9730803 | 0 | 0.01357926 | |
| GO_CYTOSOLIC_LARGE_RIBOSOMAL_SUBUNIT | | 1.7885567 | 0.00389864 | 0.03444242 | |
| GO_CYTOSOLIC_PART | | 2.1912854 | 0.00198807 | 0.00374339 | |
| GO_CYTOSOLIC_RIBOSOME | | 1.8334333 | 0.01934236 | 0.02763226 | |
| GO_CYTOSOLIC_SMALL_RIBOSOMAL_SUBUNIT | | 1.7661823 | 0.01740812 | 0.03899746 | |
| GO_DAMAGED_DNA_BINDING | | 1.8893907 | 0.00898876 | 0.01996675 | |
| GO_DEOXYRIBONUCLEASE_ACTIVITY | | 1.7681144 | 0.01094092 | 0.03875539 | |
| GO_DEOXYRIBOSE_PHOSPHATE_CATABOLIC_PROCESS | | 1.5874077 | 0.03821656 | 0.10859895 | |
| GO_DISULFIDE_OXIDOREDUCTASE_ACTIVITY | | 1.6864059 | 0.02970297 | 0.06334098 | |
| GO_DNA_BIOSYNTHETIC_PROCESS | | 1.9285553 | 0.00218818 | 0.01710591 | |
| GO_DNA_DAMAGE_RESPONSE_DETECTION_OF_DNA_DAMAGE | | 1.8489181 | 0.00847458 | 0.02583457 | |
| GO_DNA_DEPENDENT_DNA_REPLICATION | | 1.923255 | 0.01284797 | 0.01694357 | |
| GO_DNA_HELICASE_ACTIVITY | | 1.7863003 | 0.01758242 | 0.03498582 | |
| GO_DNA_POLYMERASE_BINDING | | 1.7268913 | 0.01452282 | 0.05042128 | |
| GO_DNA_REPLICATION_INITIATION | | 1.767758 | 0.01515152 | 0.0387636 | |
| GO_DNA_STRAND_ELONGATION | | 1.5571893 | 0.08351178 | 0.12483447 | |
| GO_DNA_TEMPLATED_TRANSCRIPTION_ELONGATION | | 2.2400477 | 0.00210971 | 0.00361458 | |
| GO_DNA_TEMPLATED_TRANSCRIPTION_TERMINATION | | 2.1531723 | 0.00215054 | 0.00449661 | |
| GO_ELECTRON_TRANSPORT_CHAIN | | 1.5733513 | 0.06722689 | 0.11655197 | |
| GO_ENDODEOXYRIBONUCLEASE_ACTIVITY | | 1.7145984 | 0.01511879 | 0.05419442 | |
| GO_ENDONUCLEASE_ACTIVITY | | 1.5651922 | 0.0465587 | 0.12057386 | |
| GO_ENDONUCLEASE_ACTIVITY_ACTIVE_WITH_EITHER_RIBO_OR_DEOXYRIBONUCLEIC_ACIDS_AND_PRODUCING_5_PHOSPHOMONOESTERS | | 1.933555 | 0.00420168 | 0.0167222 | |
| GO_ENDONUCLEASE_COMPLEX | | 1.9802613 | 0 | 0.01309578 | |
| GO_ENDOPEPTIDASE_COMPLEX | | 1.9471263 | 0.00206186 | 0.01571803 | |
| GO_ENDORIBONUCLEASE_ACTIVITY | | 1.5107265 | 0.05636743 | 0.15654269 | |
| GO_ENDORIBONUCLEASE_ACTIVITY_PRODUCING_5_PHOSPHOMONOESTERS | | 1.922281 | 0.00204499 | 0.01681793 | |
| GO_ENERGY_DERIVATION_BY_OXIDATION_OF_ORGANIC_COMPOUNDS | | 1.7602427 | 0.01923077 | 0.04085524 | |
| GO_ERROR_FREE_TRANSLESION_SYNTHESIS | | 1.7249316 | 0.00833333 | 0.05099789 | |
| GO_ERROR_PRONE_TRANSLESION_SYNTHESIS | | 1.8029855 | 0.01046025 | 0.03218533 | |
| GO_ESTABLISHMENT_OF_PROTEIN_LOCALIZATION_TO_CHROMOSOME | | 1.995776 | 0.00204918 | 0.01304769 | |
| GO_ESTABLISHMENT_OF_PROTEIN_LOCALIZATION_TO_ENDOPLASMIC_RETICULUM | | 1.937211 | 0.01008065 | 0.01634823 | |
| GO_ESTABLISHMENT_OF_PROTEIN_LOCALIZATION_TO_MEMBRANE | | 1.8069315 | 0.008 | 0.03155933 | |
| GO_ESTABLISHMENT_OF_PROTEIN_LOCALIZATION_TO_MITOCHONDRIAL_MEMBRANE | | 1.8276556 | 0.00206612 | 0.02812632 | |
| GO_ESTABLISHMENT_OF_PROTEIN_LOCALIZATION_TO_TELOMERE | | 1.8641248 | 0.00206186 | 0.02346929 | |
| GO_EUKARYOTIC_48S_PREINITIATION_COMPLEX | | 1.7928109 | 0 | 0.0336195 | |
| GO_EUKARYOTIC_TRANSLATION_INITIATION_FACTOR_3_COMPLEX | | 1.8314188 | 0 | 0.02768957 | |
| GO_EXODEOXYRIBONUCLEASE_ACTIVITY | | 1.6334605 | 0.0372807 | 0.08532978 | |
| GO_EXON_EXON_JUNCTION_COMPLEX | | 1.7207321 | 0.02226721 | 0.05240382 | |
| GO_EXOSOME_RNASE_COMPLEX_ | | 1.9538167 | 0.0021645 | 0.01520247 | |
| GO_FEMALE_MEIOTIC_NUCLEAR_DIVISION | | 1.592025 | 0.07157895 | 0.10645982 | |
| GO_FICOLIN_1_RICH_GRANULE | | 1.693058 | 0.04781705 | 0.0611 | |
| GO_GENERATION_OF_PRECURSOR_METABOLITES_AND_ENERGY | | 1.553867 | 0.04338843 | 0.12677354 | |
| GO_GLOBAL_GENOME_NUCLEOTIDE_EXCISION_REPAIR | | 1.950357 | 0.00860215 | 0.01544956 | |
| GO_GLYCOSYL_COMPOUND_BIOSYNTHETIC_PROCESS | | 1.6646495 | 0.02564103 | 0.07199428 | |
| GO_HEMATOPOIETIC_STEM_CELL_DIFFERENTIATION | | 1.6960298 | 0.03488372 | 0.05990372 | |
| GO_HISTONE_EXCHANGE | | 1.7735453 | 0.01742919 | 0.03780221 | |
| GO_INNER_MITOCHONDRIAL_MEMBRANE_ORGANIZATION | | 1.8540591 | 0.01101322 | 0.02516892 | |
| GO_INNER_MITOCHONDRIAL_MEMBRANE_PROTEIN_COMPLEX | | 1.9916044 | 0.00426439 | 0.01312152 | |
| GO_INTERLEUKIN_1_MEDIATED_SIGNALING_PATHWAY | | 2.0920637 | 0 | 0.00722003 | |
| GO_INTERSTRAND_CROSS_LINK_REPAIR | | 1.9147025 | 0.00220264 | 0.0174138 | |
| GO_INTRACELLULAR_PROTEIN_TRANSMEMBRANE_TRANSPORT | | 2.000829 | 0 | 0.01264928 | |
| GO_INTRAMOLECULAR_TRANSFERASE_ACTIVITY | | 1.7942084 | 0.01079914 | 0.03348633 | |
| GO_INTRINSIC_COMPONENT_OF_MITOCHONDRIAL_INNER_MEMBRANE | | 1.7368808 | 0.01118568 | 0.04738492 | |
| GO_INTRINSIC_COMPONENT_OF_MITOCHONDRIAL_MEMBRANE | | 1.6103014 | 0.05010893 | 0.09671661 | |
| GO_IRON_SULFUR_CLUSTER_ASSEMBLY | | 1.8070924 | 0.00668151 | 0.03159579 | |
| GO_ISOMERASE_ACTIVITY | | 1.5751219 | 0.02136752 | 0.11558973 | |
| GO_LARGE_RIBOSOMAL_SUBUNIT | | 2.0034301 | 0.00203252 | 0.01286257 | |
| GO_LIGASE_ACTIVITY_FORMING_CARBON_OXYGEN_BONDS | | 1.9029763 | 0.0043573 | 0.01838177 | |

| | | | |
|---|---|---|---|
| GO_MATURATION_OF_5_8S_RRNA | 1.9259416 | 0 | 0.01706409 |
| GO_MATURATION_OF_5_8S_RRNA_FROM_TRICISTRONIC_RRNA_TRANSCRIPT_SSU_RRNA_5_8S_RRNA_LSU_RRNA_ | 1.9346548 | 0 | 0.01674035 |
| GO_MATURATION_OF_LSU_RRNA | 1.9281245 | 0 | 0.01706805 |
| GO_MATURATION_OF_SSU_RRNA | 2.0549889 | 0 | 0.00900236 |
| GO_MATURATION_OF_SSU_RRNA_FROM_TRICISTRONIC_RRNA_TRANSCRIPT_SSU_RRNA_5_8S_RRNA_LSU_RRNA_ | 1.9844301 | 0 | 0.01302605 |
| GO_METAL_CLUSTER_BINDING | 1.7941837 | 0.00631579 | 0.03341325 |
| GO_METAPHASE_ANAPHASE_TRANSITION_OF_CELL_CYCLE | 1.7797961 | 0.02132196 | 0.03628285 |
| GO_METAPHASE_PLATE_CONGRESSION | 1.847305 | 0.00652174 | 0.02606935 |
| GO_MISMATCH_REPAIR | 1.4469527 | 0.08478261 | 0.20902188 |
| GO_MITOCHONDRIAL_ELECTRON_TRANSPORT_NADH_TO_UBIQUINONE | 1.6293607 | 0.06622516 | 0.08720037 |
| GO_MITOCHONDRIAL_GENE_EXPRESSION | 2.0579221 | 0 | 0.00903079 |
| GO_MITOCHONDRIAL_MATRIX | 1.8472384 | 0.01082251 | 0.02599322 |
| GO_MITOCHONDRIAL_MEMBRANE_ORGANIZATION | 1.9874864 | 0.00221239 | 0.0131214 |
| GO_MITOCHONDRIAL_MEMBRANE_PART | 1.8342129 | 0.0131579 | 0.02770652 |
| GO_MITOCHONDRIAL_PROTEIN_COMPLEX | 2.087385 | 0 | 0.00753432 |
| GO_MITOCHONDRIAL_RESPIRATORY_CHAIN_COMPLEX_ASSEMBLY | 1.9253925 | 0.00444445 | 0.01687264 |
| GO_MITOCHONDRIAL_RESPIRATORY_CHAIN_COMPLEX_I | 1.6261784 | 0.05777778 | 0.08884403 |
| GO_MITOCHONDRIAL_RNA_METABOLIC_PROCESS | 1.8401432 | 0.02542373 | 0.02691623 |
| GO_MITOCHONDRIAL_TRANSLATION | 2.04209 | 0 | 0.01072653 |
| GO_MITOCHONDRIAL_TRANSLATIONAL_TERMINATION | 2.0374904 | 0 | 0.01082413 |
| GO_MITOCHONDRIAL_TRANSMEMBRANE_TRANSPORT | 1.6178974 | 0.05010893 | 0.09333035 |
| GO_MITOCHONDRIAL_TRANSPORT | 1.8196676 | 0.01505376 | 0.02937791 |
| GO_MITOCHONDRION_ORGANIZATION | 1.9861525 | 0 | 0.01302283 |
| GO_MITOTIC_METAPHASE_PLATE_CONGRESSION | 1.8108593 | 0.00428266 | 0.03091179 |
| GO_MITOTIC_NUCLEAR_DIVISION | 1.9315755 | 0.00431965 | 0.01692344 |
| GO_MITOTIC_SISTER_CHROMATID_SEGREGATION | 1.9033178 | 0.00643777 | 0.01838307 |
| GO_MLL1_2_COMPLEX | 1.8356984 | 0.01902749 | 0.02740547 |
| GO_MRNA_5_UTR_BINDING | 2.1279755 | 0 | 0.00623267 |
| GO_MRNA_CIS_SPLICING_VIA_SPLICEOSOME | 1.5059924 | 0.11688311 | 0.15962625 |
| GO_NADH_DEHYDROGENASE_ACTIVITY | 1.7113658 | 0.02407002 | 0.05489301 |
| GO_NADH_DEHYDROGENASE_COMPLEX_ASSEMBLY | 1.8692024 | 0.00867679 | 0.02269838 |
| GO_NCRNA_3_END_PROCESSING | 2.0049105 | 0.00212766 | 0.01298959 |
| GO_NCRNA_METABOLIC_PROCESS | 2.1965227 | 0.00215983 | 0.00371978 |
| GO_NCRNA_PROCESSING | 2.2285333 | 0.0021645 | 0.00298222 |
| GO_NCRNA_TRANSCRIPTION | 2.2272177 | 0.00205339 | 0.00286752 |
| GO_NEGATIVE_REGULATION_OF_CELL_CYCLE_G2_M_PHASE_TRANSITION | 1.9621655 | 0 | 0.01464971 |
| GO_NEGATIVE_REGULATION_OF_CELL_CYCLE_PHASE_TRANSITION | 2.009199 | 0 | 0.01294595 |
| GO_NEGATIVE_REGULATION_OF_CELL_CYCLE_PROCESS | 2.0046508 | 0 | 0.01290752 |
| GO_NEGATIVE_REGULATION_OF_MITOTIC_CELL_CYCLE | 2.010031 | 0 | 0.0130065 |
| GO_NEGATIVE_REGULATION_OF_RNA_SPLICING | 1.6088134 | 0.06095238 | 0.09713763 |
| GO_NEGATIVE_REGULATION_OF_UBIQUITIN_PROTEIN_TRANSFERASE_ACTIVITY | 1.6892977 | 0.02731093 | 0.06192495 |
| GO_NF_KAPPAB_BINDING | 1.9233649 | 0 | 0.01697921 |
| GO_NUCLEAR_DNA_REPLICATION | 1.8187983 | 0.01086957 | 0.02952212 |
| GO_NUCLEAR_ENVELOPE_ORGANIZATION | 1.7314293 | 0.03501094 | 0.04904145 |
| GO_NUCLEAR_ENVELOPE_REASSEMBLY | 1.6519934 | 0.0237069 | 0.0767554 |
| GO_NUCLEAR_EXOSOME_RNASE_COMPLEX_ | 1.7324094 | 0.02335456 | 0.04883638 |
| GO_NUCLEAR_EXPORT | 2.3690681 | 0 | 0.00278036 |
| GO_NUCLEAR_TRANSCRIBED_MRNA_CATABOLIC_PROCESS | 2.2456336 | 0.00208768 | 0.0034168 |
| GO_NUCLEAR_TRANSCRIBED_MRNA_CATABOLIC_PROCESS_DEADENYLATION_DEPENDENT_DECAY | 1.7600857 | 0.02966102 | 0.04080775 |
| GO_NUCLEAR_TRANSCRIBED_MRNA_CATABOLIC_PROCESS_EXONUCLEOLYTIC | 1.9012892 | 0.00840336 | 0.01865215 |
| GO_NUCLEAR_TRANSCRIBED_MRNA_CATABOLIC_PROCESS_NONSENSE_MEDIATED_DECAY | 2.0451658 | 0.00200803 | 0.01040872 |
| GO_NUCLEAR_UBIQUITIN_LIGASE_COMPLEX | 1.8159461 | 0.01324503 | 0.02993145 |
| GO_NUCLEASE_ACTIVITY | 1.6996152 | 0.01914894 | 0.05880825 |
| GO_NUCLEIC_ACID_PHOSPHODIESTER_BOND_HYDROLYSIS | 1.872179 | 0.00867679 | 0.02212491 |
| GO_NUCLEOBASE_BIOSYNTHETIC_PROCESS | 1.7310221 | 0.01769912 | 0.04898916 |
| GO_NUCLEOBASE_CONTAINING_SMALL_MOLECULE_CATABOLIC_PROCESS | 1.4668792 | 0.03869654 | 0.19297774 |
| GO_NUCLEOID | 2.0140438 | 0 | 0.01261005 |
| GO_NUCLEOLAR_PART | 1.8013424 | 0.01077586 | 0.03253086 |
| GO_NUCLEOSIDE_MONOPHOSPHATE_BIOSYNTHETIC_PROCESS | 1.8203661 | 0.00649351 | 0.02935173 |
| GO_NUCLEOSIDE_SALVAGE | 1.6428069 | 0.0101626 | 0.08083905 |
| GO_NUCLEOSIDE_TRIPHOSPHATE_BIOSYNTHETIC_PROCESS | 1.667621 | 0.02105263 | 0.07078502 |
| GO_NUCLEOTIDE_EXCISION_REPAIR | 2.183258 | 0 | 0.0036901 |
| GO_NUCLEOTIDE_EXCISION_REPAIR_DNA_DAMAGE_RECOGNITION | 1.7783326 | 0.01054852 | 0.03649476 |
| GO_NUCLEOTIDE_EXCISION_REPAIR_DNA_DUPLEX_UNWINDING | 1.826784 | 0.00840336 | 0.02815636 |
| GO_NUCLEOTIDE_EXCISION_REPAIR_DNA_GAP_FILLING | 1.6574824 | 0.04458599 | 0.07512108 |
| GO_NUCLEOTIDE_EXCISION_REPAIR_DNA_INCISION | 2.0065653 | 0.00218818 | 0.01311337 |
| GO_NUCLEOTIDE_EXCISION_REPAIR_PREINCISION_COMPLEX_ASSEMBLY | 1.8891623 | 0.00646552 | 0.01994404 |
| GO_NUCLEOTIDE_EXCISION_REPAIR_PREINCISION_COMPLEX_STABILIZATION | 1.8392164 | 0.01072961 | 0.0270413 |
| GO_NUCLEOTIDYLTRANSFERASE_ACTIVITY | 2.0265138 | 0.0022173 | 0.01151852 |
| GO_O_METHYLTRANSFERASE_ACTIVITY | 1.439917 | 0.11316872 | 0.21418722 |
| GO_OLIGOSACCHARIDE_LIPID_INTERMEDIATE_BIOSYNTHETIC_PROCESS | 1.6417854 | 0.03586498 | 0.0810796 |
| GO_ORGANELLAR_LARGE_RIBOSOMAL_SUBUNIT | 1.9268824 | 0 | 0.01705411 |
| GO_ORGANELLAR_RIBOSOME | 2.032424 | 0 | 0.01103879 |
| GO_ORGANELLAR_SMALL_RIBOSOMAL_SUBUNIT | 1.9890507 | 0 | 0.01316798 |
| GO_ORGANELLE_ENVELOPE_LUMEN | 1.9409759 | 0.00409836 | 0.01633011 |
| GO_ORGANELLE_INNER_MEMBRANE | 1.9245344 | 0.01084599 | 0.01701663 |
| GO_OUTER_MITOCHONDRIAL_MEMBRANE_PROTEIN_COMPLEX | 1.9772155 | 0 | 0.01336573 |
| GO_OXIDATIVE_PHOSPHORYLATION | 1.8496555 | 0.01724138 | 0.02579306 |
| GO_OXIDOREDUCTASE_ACTIVITY_ACTING_ON_A_HEME_GROUP_OF_DONORS | 1.4377309 | 0.11632653 | 0.21513659 |
| GO_OXIDOREDUCTASE_ACTIVITY_ACTING_ON_NAD_P_H_QUINONE_OR_SIMILAR_COMPOUND_AS_ACCEPTOR | 1.5014703 | 0.08333334 | 0.16338961 |
| GO_OXIDOREDUCTASE_COMPLEX | 1.5858555 | 0.07484408 | 0.10967447 |
| GO_PEPTIDASE_COMPLEX | 2.0269334 | 0 | 0.01153572 |
| GO_PEPTIDYL_ARGININE_MODIFICATION | 1.4900883 | 0.06099815 | 0.1729746 |
| GO_PEPTIDYL_PROLINE_MODIFICATION | 1.6059327 | 0.05708245 | 0.09882093 |
| GO_PIGMENT_BIOSYNTHETIC_PROCESS | 1.719225 | 0.01670146 | 0.05284357 |
| GO_POLYSOMAL_RIBOSOME | 1.795513 | 0 | 0.03345429 |
| GO_POLYSOME | 2.0189724 | 0.00588235 | 0.01216475 |
| GO_POSITIVE_REGULATION_OF_CELLULAR_AMIDE_METABOLIC_PROCESS | 1.5918329 | 0.03448276 | 0.10640177 |
| GO_POSITIVE_REGULATION_OF_DNA_BIOSYNTHETIC_PROCESS | 1.5899321 | 0.04602511 | 0.10699172 |
| GO_POSITIVE_REGULATION_OF_PROTEIN_LOCALIZATION_TO_NUCLEUS | 1.7649574 | 0.01709402 | 0.03936537 |
| GO_POSITIVE_REGULATION_OF_TELOMERASE_ACTIVITY | 1.4699312 | 0.08991228 | 0.19049431 |
| GO_POSITIVE_REGULATION_OF_TELOMERASE_RNA_LOCALIZATION_TO_CAJAL_BODY | 1.7185833 | 0.00817996 | 0.05300026 |
| GO_POSITIVE_REGULATION_OF_VIRAL_PROCESS | 1.8506109 | 0.01844262 | 0.0256935 |
| GO_POSITIVE_REGULATION_OF_VIRAL_TRANSCRIPTION | 2.193418 | 0.00200401 | 0.0038437 |
| GO_POSTREPLICATION_REPAIR | 1.969971 | 0.00434783 | 0.01379843 |
| GO_PRECATALYTIC_SPLICEOSOME | 1.9453475 | 0.00420168 | 0.01580541 |

| | | | |
|---|---|---|---|
| GO_PRERIBOSOME | 2.0015123 | 0.00202429 | 0.01279714 |
| GO_PRERIBOSOME_LARGE_SUBUNIT_PRECURSOR | 1.7060698 | 0.00204082 | 0.0568016 |
| GO_PROTEASOMAL_PROTEIN_CATABOLIC_PROCESS | 1.8643802 | 0.00627615 | 0.02348525 |
| GO_PROTEASOMAL_UBIQUITIN_INDEPENDENT_PROTEIN_CATABOLIC_PROCESS | 1.5585196 | 0.05702648 | 0.1246532 |
| GO_PROTEASOME_ACCESSORY_COMPLEX | 1.7334505 | 0.00204082 | 0.04844138 |
| GO_PROTEASOME_CORE_COMPLEX | 1.6689172 | 0.00631579 | 0.07052892 |
| GO_PROTEIN_CONTAINING_COMPLEX_DISASSEMBLY | 1.9466709 | 0 | 0.01566645 |
| GO_PROTEIN_DISULFIDE_OXIDOREDUCTASE_ACTIVITY | 1.8954461 | 0.00394477 | 0.01916402 |
| GO_PROTEIN_DNA_COMPLEX_SUBUNIT_ORGANIZATION | 1.8827263 | 0.01508621 | 0.02073055 |
| GO_PROTEIN_FOLDING | 1.8376025 | 0.01242236 | 0.02712501 |
| GO_PROTEIN_IMPORT_INTO_MITOCHONDRIAL_MATRIX | 1.8602245 | 0 | 0.02398159 |
| GO_PROTEIN_INSERTION_INTO_MEMBRANE | 1.8301889 | 0.0040568 | 0.02779682 |
| GO_PROTEIN_INSERTION_INTO_MITOCHONDRIAL_MEMBRANE | 1.8315679 | 0.00632911 | 0.02773637 |
| GO_PROTEIN_K11_LINKED_UBIQUITINATION | 1.6496907 | 0.04752066 | 0.07754524 |
| GO_PROTEIN_LOCALIZATION_TO_ENDOPLASMIC_RETICULUM | 1.8916347 | 0.02235772 | 0.01975925 |
| GO_PROTEIN_LOCALIZATION_TO_MITOCHONDRION | 1.9650809 | 0.00210084 | 0.01422835 |
| GO_PROTEIN_MODIFICATION_BY_SMALL_PROTEIN_REMOVAL | 2.0329878 | 0 | 0.01114413 |
| GO_PROTEIN_NEDDYLATION | 1.7249324 | 0.01386139 | 0.05110851 |
| GO_PROTEIN_PEPTIDYL_PROLYL_ISOMERIZATION | 1.701808 | 0.04166667 | 0.05798405 |
| GO_PROTEIN_QUALITY_CONTROL_FOR_MISFOLDED_OR_INCOMPLETELY_SYNTHESIZED_PROTEINS | 1.5914953 | 0.04545455 | 0.10650186 |
| GO_PROTEIN_TARGETING | 1.9273317 | 0.0041841 | 0.01713257 |
| GO_PROTEIN_TARGETING_TO_MEMBRANE | 1.9539355 | 0.00595238 | 0.01529285 |
| GO_PROTEIN_TARGETING_TO_MITOCHONDRION | 1.9666901 | 0.00622407 | 0.01418276 |
| GO_PROTEIN_TRANSMEMBRANE_IMPORT_INTO_INTRACELLULAR_ORGANELLE | 1.9711045 | 0 | 0.01373525 |
| GO_PROTEIN_TRANSMEMBRANE_TRANSPORT | 1.9748048 | 0.00205339 | 0.01340614 |
| GO_PROTEIN_TRANSPORTER_ACTIVITY | 1.6587024 | 0.0256917 | 0.07462262 |
| GO_PROTON_TRANSPORTING_TWO_SECTOR_ATPASE_COMPLEX | 1.5926589 | 0.0421941 | 0.10629695 |
| GO_PSEUDOURIDINE_SYNTHESIS | 1.896903 | 0.00212766 | 0.01897282 |
| GO_PURINE_NUCLEOSIDE_BIOSYNTHETIC_PROCESS | 1.4276378 | 0.08092485 | 0.2254352 |
| GO_PURINE_NUCLEOSIDE_MONOPHOSPHATE_BIOSYNTHETIC_PROCESS | 1.8114953 | 0 | 0.03078106 |
| GO_PYRIMIDINE_CONTAINING_COMPOUND_BIOSYNTHETIC_PROCESS | 1.4802861 | 0.05781585 | 0.1813439 |
| GO_PYRIMIDINE_NUCLEOSIDE_MONOPHOSPHATE_BIOSYNTHETIC_PROCESS | 1.6107582 | 0.04250559 | 0.09690166 |
| GO_PYRIMIDINE_NUCLEOTIDE_BIOSYNTHETIC_PROCESS | 1.5574156 | 0.03303965 | 0.12483344 |
| GO_PYRIMIDINE_RIBONUCLEOTIDE_BIOSYNTHETIC_PROCESS | 1.4740709 | 0.07272727 | 0.1869442 |
| GO_REGULATION_OF_CELL_CYCLE_ARREST | 1.6915 | 0.01059322 | 0.06140494 |
| GO_REGULATION_OF_CELL_CYCLE_G2_M_PHASE_TRANSITION | 1.9057531 | 0.01121076 | 0.01801452 |
| GO_REGULATION_OF_CELL_CYCLE_PHASE_TRANSITION | 1.9901812 | 0.004329 | 0.01318346 |
| GO_REGULATION_OF_CELLULAR_AMIDE_METABOLIC_PROCESS | 1.8395616 | 0.0021645 | 0.02702067 |
| GO_REGULATION_OF_CHROMOSOME_SEPARATION | 1.7771627 | 0.02547771 | 0.03679544 |
| GO_REGULATION_OF_DNA_BIOSYNTHETIC_PROCESS | 1.6565903 | 0.01793722 | 0.07525609 |
| GO_REGULATION_OF_DNA_TEMPLATED_TRANSCRIPTION_IN_RESPONSE_TO_STRESS | 1.8229514 | 0.01666667 | 0.02880438 |
| GO_REGULATION_OF_ESTABLISHMENT_OF_PLANAR_POLARITY | 1.5236623 | 0.07113821 | 0.14769147 |
| GO_REGULATION_OF_HEMATOPOIETIC_PROGENITOR_CELL_DIFFERENTIATION | 1.9520475 | 0.00203252 | 0.01535581 |
| GO_REGULATION_OF_MITOCHONDRIAL_OUTER_MEMBRANE_PERMEABILIZATION_INVOLVED_IN_APOPTOTIC_SIGNALING_PATHWAY | 1.7578232 | 0.01716738 | 0.04127476 |
| GO_REGULATION_OF_MITOCHONDRIAL_TRANSLATION | 1.9585555 | 0 | 0.01473789 |
| GO_REGULATION_OF_MRNA_CATABOLIC_PROCESS | 1.8051983 | 0.01483051 | 0.03184892 |
| GO_REGULATION_OF_RELEASE_OF_CYTOCHROME_C_FROM_MITOCHONDRIA | 1.4047052 | 0.04862579 | 0.24903302 |
| GO_REGULATION_OF_STEM_CELL_DIFFERENTIATION | 1.498783 | 0.10060363 | 0.16538422 |
| GO_REGULATION_OF_TELOMERASE_ACTIVITY | 1.5337137 | 0.05111111 | 0.14033787 |
| GO_REGULATION_OF_TRANSCRIPTION_FROM_RNA_POLYMERASE_II_PROMOTER_IN_RESPONSE_TO_HYPOXIA | 2.0926378 | 0 | 0.00733306 |
| GO_REGULATION_OF_TRANSLATIONAL_FIDELITY | 1.6982397 | 0.03043478 | 0.05923051 |
| GO_REGULATION_OF_TRANSLATIONAL_INITIATION | 2.087771 | 0 | 0.00755925 |
| GO_REGULATION_OF_UBIQUITIN_PROTEIN_LIGASE_ACTIVITY | 1.6895868 | 0.02401747 | 0.06201919 |
| GO_REGULATION_OF_VIRAL_TRANSCRIPTION | 1.8461814 | 0.00806452 | 0.02617354 |
| GO_RELEASE_OF_CYTOCHROME_C_FROM_MITOCHONDRIA | 1.515947 | 0.03144654 | 0.15263557 |
| GO_REPLICATION_FORK | 1.8122905 | 0.02212389 | 0.03094333 |
| GO_REPLISOME | 1.7041947 | 0.04595186 | 0.05727156 |
| GO_RESPIRASOME | 1.8450825 | 0.01282051 | 0.02624841 |
| GO_RESPIRATORY_CHAIN_COMPLEX | 1.8119128 | 0.01528384 | 0.03085884 |
| GO_RESPIRATORY_ELECTRON_TRANSPORT_CHAIN | 1.8547392 | 0.01766004 | 0.02506167 |
| GO_RIBONUCLEASE_ACTIVITY | 1.528741 | 0.05052632 | 0.14467703 |
| GO_RIBONUCLEOPROTEIN_COMPLEX_BINDING | 2.1829453 | 0 | 0.00359299 |
| GO_RIBONUCLEOPROTEIN_COMPLEX_BIOGENESIS | 2.342016 | 0 | 0.00246015 |
| GO_RIBONUCLEOPROTEIN_COMPLEX_SUBUNIT_ORGANIZATION | 2.3244767 | 0.00210084 | 0.00224505 |
| GO_RIBONUCLEOSIDE_MONOPHOSPHATE_BIOSYNTHETIC_PROCESS | 1.9671965 | 0 | 0.01422098 |
| GO_RIBONUCLEOSIDE_TRIPHOSPHATE_BIOSYNTHETIC_PROCESS | 1.6747597 | 0.0375 | 0.0684026 |
| GO_RIBOSOMAL_LARGE_SUBUNIT_ASSEMBLY | 1.9034191 | 0 | 0.01844522 |
| GO_RIBOSOMAL_LARGE_SUBUNIT_BIOGENESIS | 2.065284 | 0 | 0.00828253 |
| GO_RIBOSOMAL_SMALL_SUBUNIT_ASSEMBLY | 1.8340234 | 0.00403226 | 0.02764836 |
| GO_RIBOSOMAL_SMALL_SUBUNIT_BIOGENESIS | 2.1569118 | 0 | 0.00436729 |
| GO_RIBOSOMAL_SUBUNIT | 2.0386953 | 0.00201207 | 0.01100966 |
| GO_RIBOSOME | 2.0322864 | 0.00613497 | 0.01093558 |
| GO_RIBOSOME_ASSEMBLY | 2.1010652 | 0 | 0.00741616 |
| GO_RIBOSOME_BINDING | 2.00544 | 0 | 0.01306835 |
| GO_RIBOSOME_BIOGENESIS | 2.2326105 | 0 | 0.00320903 |
| GO_RNA_3_END_PROCESSING | 2.4100187 | 0 | 0.00530343 |
| GO_RNA_5_END_PROCESSING | 1.6780379 | 0.02760085 | 0.06712114 |
| GO_RNA_CAP_BINDING | 1.9230462 | 0.00401606 | 0.01686891 |
| GO_RNA_CAPPING | 2.1838481 | 0.002079 | 0.00372528 |
| GO_RNA_CATABOLIC_PROCESS | 2.2558725 | 0 | 0.00346842 |
| GO_RNA_DEPENDENT_DNA_BIOSYNTHETIC_PROCESS | 2.2570162 | 0 | 0.00357241 |
| GO_RNA_EXPORT_FROM_NUCLEUS | 2.3827388 | 0.00223214 | 0.00463309 |
| GO_RNA_METHYLATION | 1.7143462 | 0.02540416 | 0.05418176 |
| GO_RNA_METHYLTRANSFERASE_ACTIVITY | 1.805025 | 0.02022472 | 0.03184271 |
| GO_RNA_MODIFICATION | 1.900166 | 0.00451467 | 0.01866726 |
| GO_RNA_PHOSPHODIESTER_BOND_HYDROLYSIS | 1.7295674 | 0.02536998 | 0.04956748 |
| GO_RNA_PHOSPHODIESTER_BOND_HYDROLYSIS_ENDONUCLEOLYTIC | 1.5137452 | 0.06105263 | 0.15431039 |
| GO_RNA_POLYMERASE_ACTIVITY | 2.0828018 | 0 | 0.00780415 |
| GO_RNA_POLYMERASE_COMPLEX | 2.3010607 | 0 | 0.00283624 |
| GO_RNA_POLYMERASE_II_CORE_COMPLEX | 1.7821238 | 0.01214575 | 0.03590388 |
| GO_RNA_POLYMERASE_II_HOLOENZYME | 2.3081698 | 0 | 0.0029891 |
| GO_RNA_POLYMERASE_III_ACTIVITY | 1.9981948 | 0.00211417 | 0.01284149 |
| GO_RNA_POLYMERASE_III_COMPLEX | 1.9933542 | 0 | 0.01314387 |
| GO_RNA_SPLICING | 2.2385137 | 0.00450451 | 0.00325312 |
| GO_RNA_SPLICING_VIA_TRANSESTERIFICATION_REACTIONS | 2.265842 | 0.00445434 | 0.00360919 |
| GO_ROUGH_ENDOPLASMIC_RETICULUM | 1.4393644 | 0.04511278 | 0.21404195 |

| | | | |
|---|---|---|---|
| GO_ROUGH_ENDOPLASMIC_RETICULUM_MEMBRANE | 1.4561732 | 0.05996132 | 0.20154174 |
| GO_RRNA_BINDING | 2.074625 | 0.00207039 | 0.00803262 |
| GO_RRNA_CONTAINING_RIBONUCLEOPROTEIN_COMPLEX_EXPORT_FROM_NUCLEUS | 1.7955228 | 0 | 0.03354594 |
| GO_RRNA_METABOLIC_PROCESS | 2.2296433 | 0 | 0.00310648 |
| GO_RRNA_METHYLATION | 1.6429317 | 0.02721088 | 0.08089889 |
| GO_RRNA_MODIFICATION | 1.8268757 | 0.01382489 | 0.02822601 |
| GO_RRNA_TRANSCRIPTION | 1.7976512 | 0.02004008 | 0.03318499 |
| GO_S_ADENOSYLMETHIONINE_DEPENDENT_METHYLTRANSFERASE_ACTIVITY | 1.6776258 | 0.02863436 | 0.06722403 |
| GO_SCF_DEPENDENT_PROTEASOMAL_UBIQUITIN_DEPENDENT_PROTEIN_CATABOLIC_PROCESS | 2.0352366 | 0.00205339 | 0.01094633 |
| GO_SINGLE_STRANDED_DNA_BINDING | 2.1147523 | 0 | 0.00652656 |
| GO_SM_LIKE_PROTEIN_FAMILY_COMPLEX | 2.1104994 | 0 | 0.00679274 |
| GO_SMALL_NUCLEOLAR_RIBONUCLEOPROTEIN_COMPLEX | 2.0710263 | 0 | 0.00821773 |
| GO_SMALL_RIBOSOMAL_SUBUNIT | 1.9799579 | 0.00398406 | 0.01308186 |
| GO_SMALL_SUBUNIT_PROCESSOME | 1.8875377 | 0.00809717 | 0.02011306 |
| GO_SMN_SM_PROTEIN_COMPLEX | 1.8915982 | 0.00197239 | 0.01968319 |
| GO_SNORNA_BINDING | 1.9500047 | 0 | 0.01540676 |
| GO_SNRNA_3_END_PROCESSING | 1.7757419 | 0.02145923 | 0.03725673 |
| GO_SNRNA_BINDING | 1.8933254 | 0.00838574 | 0.01952337 |
| GO_SNRNA_PROCESSING | 1.7720416 | 0.02202643 | 0.03775061 |
| GO_SPLICEOSOMAL_COMPLEX | 2.1985698 | 0.00225734 | 0.00382255 |
| GO_SPLICEOSOMAL_SNRNP_ASSEMBLY | 1.9224453 | 0.00614754 | 0.01686125 |
| GO_SPLICEOSOMAL_TRI_SNRNP_COMPLEX | 1.9164953 | 0.00419287 | 0.01730064 |
| GO_STRUCTURAL_CONSTITUENT_OF_RIBOSOME | 1.9983335 | 0.00205339 | 0.01294077 |
| GO_TELOMERASE_HOLOENZYME_COMPLEX | 1.9196061 | 0 | 0.01693326 |
| GO_TELOMERASE_RNA_BINDING | 1.8115909 | 0.00643777 | 0.0308517 |
| GO_TELOMERASE_RNA_LOCALIZATION | 1.7837712 | 0.0101833 | 0.03553075 |
| GO_TELOMERE_MAINTENANCE_VIA_SEMI_CONSERVATIVE_REPLICATION | 1.8012161 | 0.00643777 | 0.03245752 |
| GO_TELOMERE_MAINTENANCE_VIA_TELOMERE_LENGTHENING | 2.239983 | 0 | 0.00342434 |
| GO_TELOMERE_ORGANIZATION | 2.1649547 | 0 | 0.00434309 |
| GO_TERMINATION_OF_RNA_POLYMERASE_I_TRANSCRIPTION | 1.9154474 | 0.00652174 | 0.01738848 |
| GO_TERMINATION_OF_RNA_POLYMERASE_II_TRANSCRIPTION | 1.9812268 | 0.00617284 | 0.0130393 |
| GO_TETRAPYRROLE_BIOSYNTHETIC_PROCESS | 1.4077245 | 0.16075157 | 0.24592766 |
| GO_THREONINE_TYPE_PEPTIDASE_ACTIVITY | 1.6828439 | 0.01048218 | 0.06504586 |
| GO_TRANSCRIPTION_BY_RNA_POLYMERASE_I | 2.0015345 | 0.00860215 | 0.01291513 |
| GO_TRANSCRIPTION_BY_RNA_POLYMERASE_III | 1.8672941 | 0.01035197 | 0.02297067 |
| GO_TRANSCRIPTION_COUPLED_NUCLEOTIDE_EXCISION_REPAIR | 2.1266692 | 0 | 0.00617799 |
| GO_TRANSCRIPTION_ELONGATION_FROM_RNA_POLYMERASE_I_PROMOTER | 1.9243308 | 0.00668151 | 0.01696481 |
| GO_TRANSCRIPTION_ELONGATION_FROM_RNA_POLYMERASE_II_PROMOTER | 2.2818427 | 0 | 0.00306753 |
| GO_TRANSCRIPTION_INITIATION_FROM_RNA_POLYMERASE_I_PROMOTER | 1.8296685 | 0.00898876 | 0.02784528 |
| GO_TRANSCRIPTION_PREINITIATION_COMPLEX_ASSEMBLY | 1.7708912 | 0.0251046 | 0.03806444 |
| GO_TRANSFERASE_ACTIVITY_TRANSFERRING_ONE_CARBON_GROUPS | 1.5627588 | 0.04068523 | 0.122158 |
| GO_TRANSLATION_ELONGATION_FACTOR_ACTIVITY | 1.6362388 | 0.02906977 | 0.08406939 |
| GO_TRANSLATION_FACTOR_ACTIVITY_RNA_BINDING | 2.0785854 | 0.00401606 | 0.00795049 |
| GO_TRANSLATION_INITIATION_FACTOR_ACTIVITY | 2.1172335 | 0 | 0.0066128 |
| GO_TRANSLATION_INITIATION_FACTOR_BINDING | 1.6316019 | 0.03612167 | 0.08627068 |
| GO_TRANSLATION_PREINITIATION_COMPLEX | 1.7974648 | 0 | 0.03315038 |
| GO_TRANSLATION_REGULATOR_ACTIVITY | 2.017788 | 0.00201613 | 0.01230293 |
| GO_TRANSLATION_REGULATOR_ACTIVITY_NUCLEIC_ACID_BINDING | 1.9776208 | 0.0020202 | 0.01341094 |
| GO_TRANSLATIONAL_ELONGATION | 2.159396 | 0 | 0.00438025 |
| GO_TRANSLATIONAL_INITIATION | 2.1869361 | 0.00207039 | 0.00359186 |
| GO_TRANSLATIONAL_TERMINATION | 2.1199782 | 0 | 0.00652231 |
| GO_TRANSLESION_SYNTHESIS | 1.9241294 | 0.00220751 | 0.01690564 |
| GO_TRICARBOXYLIC_ACID_CYCLE | 1.5224681 | 0.12184874 | 0.14799733 |
| GO_TRNA_5_END_PROCESSING | 1.7466893 | 0.00425532 | 0.04470521 |
| GO_TRNA_BINDING | 1.9615257 | 0.00824742 | 0.01464234 |
| GO_TRNA_METABOLIC_PROCESS | 2.0974853 | 0.00438597 | 0.00728619 |
| GO_TRNA_METHYLATION | 1.5490136 | 0.08278867 | 0.12976718 |
| GO_TRNA_MODIFICATION | 1.9141709 | 0.01079914 | 0.01740481 |
| GO_TRNA_PROCESSING | 2.0039368 | 0.00894855 | 0.01290552 |
| GO_TRNA_SPECIFIC_RIBONUCLEASE_ACTIVITY | 1.7694962 | 0.01079914 | 0.0384407 |
| GO_TRNA_WOBBLE_BASE_MODIFICATION | 1.569126 | 0.0371134 | 0.11822939 |
| GO_U1_SNRNP | 1.5857756 | 0.05776893 | 0.10958181 |
| GO_U12_TYPE_SPLICEOSOMAL_COMPLEX | 1.9263049 | 0.0021322 | 0.01709877 |
| GO_U2_SNRNP | 1.939954 | 0 | 0.01634624 |
| GO_U2_TYPE_CATALYTIC_STEP_2_SPLICEOSOME | 1.9381659 | 0 | 0.01637241 |
| GO_U2_TYPE_SPLICEOSOMAL_COMPLEX | 2.0664234 | 0.00436681 | 0.00841384 |
| GO_U5_SNRNP | 1.788486 | 0.01642711 | 0.03436942 |
| GO_UBIQUITIN_PROTEIN_TRANSFERASE_REGULATOR_ACTIVITY | 1.7168332 | 0.0041841 | 0.05340308 |
| GO_UNFOLDED_PROTEIN_BINDING | 1.9328228 | 0.00618557 | 0.01675258 |
| GO_VIRAL_GENE_EXPRESSION | 2.235821 | 0 | 0.00321955 |
| HALLMARK_DNA_REPAIR | 2.2431881 | 0 | 0 |
| HALLMARK_E2F_TARGETS | 1.8357594 | 0 | 0.0158998 |
| HALLMARK_MTORC1_SIGNALING | 1.948551 | 0.00211417 | 0.00697068 |
| HALLMARK_MYC_TARGETS_V1 | 2.0653164 | 0 | 0.00245984 |
| HALLMARK_MYC_TARGETS_V2 | 1.9752315 | 0 | 0.00638954 |
| HALLMARK_OXIDATIVE_PHOSPHORYLATION | 1.8664356 | 0.00970874 | 0.01333482 |
| HALLMARK_UNFOLDED_PROTEIN_RESPONSE | 2.0384197 | 0 | 0.00302806 |
| HALLMARK_UV_RESPONSE_UP | 1.472364 | 0.05060729 | 0.19898874 |
| KEGG_ALZHEIMERS_DISEASE | 1.5431873 | 0.086 | 0.17683585 |
| KEGG_AMINOACYL_TRNA_BIOSYNTHESIS | 1.882898 | 0.00829876 | 0.02469181 |
| KEGG_BASE_EXCISION_REPAIR | 1.6271341 | 0.04989605 | 0.11365747 |
| KEGG_DNA_REPLICATION | 1.6245027 | 0.03571429 | 0.11010298 |
| KEGG_HOMOLOGOUS_RECOMBINATION | 1.649009 | 0.0260521 | 0.10039916 |
| KEGG_HUNTINGTONS_DISEASE | 1.8896232 | 0.01419878 | 0.03001381 |
| KEGG_MISMATCH_REPAIR | 1.7011684 | 0.0125 | 0.07942785 |
| KEGG_NUCLEOTIDE_EXCISION_REPAIR | 1.877586 | 0.01473684 | 0.02341295 |
| KEGG_ONE_CARBON_POOL_BY_FOLATE | 1.4823636 | 0.07784431 | 0.22015722 |
| KEGG_OXIDATIVE_PHOSPHORYLATION | 1.8853046 | 0.02857143 | 0.02762242 |
| KEGG_PARKINSONS_DISEASE | 1.6665127 | 0.07480315 | 0.09438139 |
| KEGG_PROTEASOME | 1.7589918 | 0 | 0.04971591 |
| KEGG_PROTEIN_EXPORT | 1.6999947 | 0.02941177 | 0.07536143 |
| KEGG_PYRIMIDINE_METABOLISM | 1.9321352 | 0.00409836 | 0.01964345 |
| KEGG_RIBOSOME | 1.8032627 | 0.00369686 | 0.03861523 |
| KEGG_RNA_DEGRADATION | 2.096173 | 0.00211417 | 0.01113524 |
| KEGG_RNA_POLYMERASE | 1.976757 | 0.00209205 | 0.01972652 |
| KEGG_SPLICEOSOME | 2.1573892 | 0.00636943 | 0.0102908 |

| | | | |
|---|---|---|---|
| MUHAR_MYC | 2.056677 | 0 | 0 |
| REACTOME_ABC_TRANSPORTER_DISORDERS | 1.5155689 | 0.06144068 | 0.11790481 |
| REACTOME_ABORTIVE_ELONGATION_OF_HIV_1_TRANSCRIPT_IN_THE_ABSENCE_OF_TAT | 2.0941167 | 0 | 0.00455345 |
| REACTOME_ACTIVATION_OF_APC_C_AND_APC_C:CDC20_MEDIATED_DEGRADATION_OF_MITOTIC_PROTEINS | 1.8943316 | 0 | 0.0140302 |
| REACTOME_ACTIVATION_OF_ATR_IN_RESPONSE_TO_REPLICATION_STRESS | 1.7317466 | 0.00674157 | 0.03799924 |
| REACTOME_ACTIVATION_OF_NF_KAPPAB_IN_B_CELLS | 1.7859186 | 0.01649485 | 0.02643907 |
| REACTOME_ACTIVATION_OF_THE_MRNA_UPON_BINDING_OF_THE_CAP_BINDING_COMPLEX_AND_EIFS_AND_SUBSEQUENT_BINDING_TO_43S | 1.9022554 | 0.00201613 | 0.01364553 |
| REACTOME_ACTIVATION_OF_THE_PRE_REPLICATIVE_COMPLEX | 1.6589166 | 0.01565995 | 0.05832152 |
| REACTOME_ANTIGEN_PROCESSING_CROSS_PRESENTATION | 1.5519149 | 0.08146639 | 0.09980982 |
| REACTOME_ANTIGEN_PROCESSING:_UBIQUITINATION_PROTEASOME_DEGRADATION | 1.84013 | 0.00700935 | 0.01891318 |
| REACTOME_APC_C:CDC20_MEDIATED_DEGRADATION_OF_CYCLIN_B | 1.8337784 | 0 | 0.01981006 |
| REACTOME_APC_C:CDH1_MEDIATED_DEGRADATION_OF_CDC20_AND_OTHER_APC_C:CDH1_TARGETED_PROTEINS_IN_LATE_MITOSIS_EARLY_G1 | 1.9559425 | 0 | 0.00965798 |
| REACTOME_APC_CDC20_MEDIATED_DEGRADATION_OF_NEK2A | 1.849929 | 0 | 0.018104 |
| REACTOME_ASSEMBLY_OF_THE_PRE_REPLICATIVE_COMPLEX | 1.8740636 | 0.0021692 | 0.01550704 |
| REACTOME_ASSOCIATION_OF_TRIC_CCT_WITH_TARGET_PROTEINS_DURING_BIOSYNTHESIS | 1.804524 | 0.01535088 | 0.02352658 |
| REACTOME_ASYMMETRIC_LOCALIZATION_OF_PCP_PROTEINS | 1.7341 | 0.04928131 | 0.03790209 |
| REACTOME_ATF4_ACTIVATES_GENES_IN_RESPONSE_TO_ENDOPLASMIC_RETICULUM_STRESS | 1.7504379 | 0.02857143 | 0.03422477 |
| REACTOME_AUF1_HNRNP_D0_BINDS_AND_DESTABILIZES_MRNA | 1.8840169 | 0 | 0.01498933 |
| REACTOME_AUTOPHAGY | 1.568769 | 0.05579399 | 0.0925246 |
| REACTOME_BASE_EXCISION_REPAIR | 1.7733467 | 0.01360544 | 0.02916792 |
| REACTOME_BUDDING_AND_MATURATION_OF_HIV_VIRION | 1.333582 | 0.17805383 | 0.23225749 |
| REACTOME_BUTYRATE_RESPONSE_FACTOR_1_BRF1_BINDS_AND_DESTABILIZES_MRNA | 1.5610219 | 0.07727273 | 0.09639601 |
| REACTOME_C_TYPE_LECTIN_RECEPTORS_CLRS | 1.345016 | 0.16044776 | 0.22483985 |
| REACTOME_CDK_MEDIATED_PHOSPHORYLATION_AND_REMOVAL_OF_CDC6 | 2.0102255 | 0 | 0.00648616 |
| REACTOME_CELL_CYCLE_CHECKPOINTS | 1.9064995 | 0.00223714 | 0.01350112 |
| REACTOME_CELL_CYCLE_MITOTIC | 1.9226395 | 0.00451467 | 0.01226561 |
| REACTOME_CELLULAR_RESPONSE_TO_HYPOXIA | 2.038795 | 0.00210971 | 0.00526397 |
| REACTOME_CELLULAR_RESPONSES_TO_EXTERNAL_STIMULI | 2.0772338 | 0 | 0.00471376 |
| REACTOME_CELLULAR_RESPONSES_TO_STRESS | 2.0289843 | 0.0021692 | 0.00533264 |
| REACTOME_CHROMOSOME_MAINTENANCE | 1.779686 | 0.00450451 | 0.02760413 |
| REACTOME_CITRIC_ACID_CYCLE_TCA_CYCLE | 1.685309 | 0.04237288 | 0.05091537 |
| REACTOME_CLEC7A_DECTIN_1_SIGNALING | 1.866092 | 0.00626305 | 0.01640476 |
| REACTOME_CLEC7A_DECTIN_1_SIGNALING | 1.866092 | 0.00626305 | 0.01640476 |
| REACTOME_COMPLEX_I_BIOGENESIS | 1.8396434 | 0.01545254 | 0.01889371 |
| REACTOME_COOPERATION_OF_PREFOLDIN_AND_TRIC_CCT_IN_ACTIN_AND_TUBULIN_FOLDING | 1.8571782 | 0.00413223 | 0.01725428 |
| REACTOME_CROSS_PRESENTATION_OF_SOLUBLE_EXOGENOUS_ANTIGENS_ENDOSOMES | 1.7670207 | 0.00650759 | 0.03007402 |
| REACTOME_CYCLIN_A_B1_B2_ASSOCIATED_EVENTS_DURING_G2_M_TRANSITION | 1.5748835 | 0.04625551 | 0.09042875 |
| REACTOME_CYCLIN_A:CDK2_ASSOCIATED_EVENTS_AT_S_PHASE_ENTRY | 2.0822902 | 0 | 0.00469404 |
| REACTOME_CYTOSOLIC_SENSORS_OF_PATHOGEN_ASSOCIATED_DNA | 1.8536217 | 0.00652174 | 0.01775036 |
| REACTOME_CYTOSOLIC_TRNA_AMINOACYLATION | 1.714602 | 0.00425532 | 0.04235866 |
| REACTOME_DEADENYLATION_DEPENDENT_MRNA_DECAY | 1.9930372 | 0.00689655 | 0.00719349 |
| REACTOME_DECTIN_1_MEDIATED_NONCANONICAL_NF_KB_SIGNALING | 1.9134146 | 0.0040568 | 0.01277288 |
| REACTOME_DEFECTIVE_CFTR_CAUSES_CYSTIC_FIBROSIS | 1.8812842 | 0.00819672 | 0.01507027 |
| REACTOME_DEGRADATION_OF_AXIN | 1.7047421 | 0.01914894 | 0.04568336 |
| REACTOME_DEGRADATION_OF_BETA_CATENIN_BY_THE_DESTRUCTION_COMPLEX | 1.7323086 | 0.03893443 | 0.03799569 |
| REACTOME_DEGRADATION_OF_DVL | 1.9369954 | 0 | 0.0109852 |
| REACTOME_DEGRADATION_OF_GLI1_BY_THE_PROTEASOME | 1.8235619 | 0.00208768 | 0.02138371 |
| REACTOME_DISEASES_ASSOCIATED_WITH_N_GLYCOSYLATION_OF_PROTEINS | 1.6708344 | 0.02826087 | 0.05465856 |
| REACTOME_DNA_DAMAGE_BYPASS | 1.8984023 | 0.01133787 | 0.0138251 |
| REACTOME_DNA_DAMAGE_RECOGNITION_IN_GG_NER | 2.0199854 | 0.00453515 | 0.00587098 |
| REACTOME_DNA_REPAIR | 2.0679429 | 0.002331 | 0.00488687 |
| REACTOME_DNA_REPLICATION | 1.8128762 | 0.00456621 | 0.02275313 |
| REACTOME_DNA_REPLICATION_PRE_INITIATION | 1.8186159 | 0.00449438 | 0.0218719 |
| REACTOME_DNA_STRAND_ELONGATION | 1.58576 | 0.02721088 | 0.08620298 |
| REACTOME_DOWNSTREAM_SIGNALING_EVENTS_OF_B_CELL_RECEPTOR_BCR | 1.7936503 | 0.02263374 | 0.02515944 |
| REACTOME_DOWNSTREAM_TCR_SIGNALING | 1.4156517 | 0.18181819 | 0.1748493 |
| REACTOME_DUAL_INCISION_IN_GG_NER | 1.8893409 | 0.00681818 | 0.01453917 |
| REACTOME_DUAL_INCISION_IN_TC_NER | 2.0519087 | 0 | 0.00524419 |
| REACTOME_E2F_MEDIATED_REGULATION_OF_DNA_REPLICATION | 1.4963439 | 0.12211981 | 0.12783322 |
| REACTOME_ENDOSOMAL_SORTING_COMPLEX_REQUIRED_FOR_TRANSPORT_ESCRT | 1.402968 | 0.13168724 | 0.18276492 |
| REACTOME_EUKARYOTIC_TRANSLATION_INITIATION | 1.8935987 | 0.00205761 | 0.01400914 |
| REACTOME_EXTENSION_OF_TELOMERES | 1.6693677 | 0.01830664 | 0.05512565 |
| REACTOME_FANCONI_ANEMIA_PATHWAY | 1.8858042 | 0.00444445 | 0.01493684 |
| REACTOME_FBXL7_DOWN_REGULATES_AURKA_DURING_MITOTIC_ENTRY_AND_IN_EARLY_MITOSIS | 1.884426 | 0 | 0.01510691 |
| REACTOME_FGFR2_ALTERNATIVE_SPLICING | 2.0705 | 0.00429185 | 0.00499285 |
| REACTOME_FORMATION_OF_HIV_ELONGATION_COMPLEX_IN_THE_ABSENCE_OF_HIV_TAT | 2.2766817 | 0 | 9.08E-04 |
| REACTOME_FORMATION_OF_INCISION_COMPLEX_IN_GG_NER | 2.0931168 | 0.00223714 | 0.00430732 |
| REACTOME_FORMATION_OF_TC_NER_PRE_INCISION_COMPLEX | 2.166558 | 0 | 0.00312782 |
| REACTOME_FORMATION_OF_THE_EARLY_ELONGATION_COMPLEX | 2.1862793 | 0 | 0.00212476 |
| REACTOME_FORMATION_OF_TUBULIN_FOLDING_INTERMEDIATES_BY_CCT_TRIC | 1.7275188 | 0.02385686 | 0.03879828 |
| REACTOME_G1_PHASE | 1.6747853 | 0.01580136 | 0.05385908 |
| REACTOME_G1_S_DNA_DAMAGE_CHECKPOINTS | 1.969583 | 0 | 0.00896889 |
| REACTOME_G2_M_CHECKPOINTS | 1.86807 | 0.00453515 | 0.01626517 |
| REACTOME_GAP_FILLING_DNA_REPAIR_SYNTHESIS_AND_LIGATION_IN_GG_NER | 1.6202799 | 0.05287356 | 0.07226455 |
| REACTOME_GENE_AND_PROTEIN_EXPRESSION_BY_JAK_STAT_SIGNALING_AFTER_INTERLEUKIN_12_STIMULATION | 1.7681276 | 0.01082251 | 0.02998463 |
| REACTOME_GLOBAL_GENOME_NUCLEOTIDE_EXCISION_REPAIR_GG_NER | 2.047779 | 0.00234742 | 0.00543588 |
| REACTOME_GLUCOSE_METABOLISM | 1.6745337 | 0.02783726 | 0.05376211 |
| REACTOME_HDR_THROUGH_HOMOLOGOUS_RECOMBINATION_HRR | 1.8148118 | 0.02192983 | 0.02256042 |
| REACTOME_HEDGEHOG_LIGAND_BIOGENESIS | 1.6083832 | 0.06126482 | 0.07701983 |
| REACTOME_HEDGEHOG_OFF_STATE | 1.3765708 | 0.14718615 | 0.20122658 |
| REACTOME_HIV_ELONGATION_ARREST_AND_RECOVERY | 2.1582744 | 0 | 0.00320618 |
| REACTOME_HIV_INFECTION | 2.242397 | 0 | 0.00164362 |
| REACTOME_HIV_LIFE_CYCLE | 2.3258238 | 0 | 0.00107082 |
| REACTOME_HOST_INTERACTIONS_OF_HIV_FACTORS | 1.9976398 | 0.00212314 | 0.00714514 |
| REACTOME_HSF1_ACTIVATION | 1.5403883 | 0.09421842 | 0.10544926 |
| REACTOME_INFECTIOUS_DISEASE | 2.3802943 | 0 | 0.00308723 |
| REACTOME_INFLUENZA_INFECTION | 2.1646502 | 0 | 0.00307655 |
| REACTOME_INHIBITION_OF_THE_PROTEOLYTIC_ACTIVITY_OF_APC_C_REQUIRED_FOR_THE_ONSET_OF_ANAPHASE_BY_MITOTIC_SPINDLE_CHECKPOINT_COMPONENTS | 1.7867084 | 0 | 0.02638664 |
| REACTOME_INSERTION_OF_TAIL_ANCHORED_PROTEINS_INTO_THE_ENDOPLASMIC_RETICULUM_MEMBRANE | 1.6032367 | 0.04938272 | 0.07877643 |
| REACTOME_INTERACTIONS_OF_REV_WITH_HOST_CELLULAR_PROTEINS | 2.0082417 | 0 | 0.00650114 |
| REACTOME_INTERACTIONS_OF_VPR_WITH_HOST_CELLULAR_PROTEINS | 2.0919504 | 0 | 0.00429362 |
| REACTOME_INTERCONVERSION_OF_NUCLEOTIDE_DI_AND_TRIPHOSPHATES | 1.5195963 | 0.06029106 | 0.11589322 |
| REACTOME_INTERLEUKIN_1_FAMILY_SIGNALING | 1.8979396 | 0.00622407 | 0.01374058 |
| REACTOME_INTERLEUKIN_1_SIGNALING | 2.0371792 | 0 | 0.00522422 |
| REACTOME_KSRP_KHSRP_BINDS_AND_DESTABILIZES_MRNA | 1.7661648 | 0.01431981 | 0.03019848 |
| REACTOME_LAGGING_STRAND_SYNTHESIS | 1.5559177 | 0.05263158 | 0.09838529 |

| | | | |
|---|---|---|---|
| REACTOME_M_PHASE | 1.988349 | 0.00223714 | 0.00746962 |
| REACTOME_MACROAUTOPHAGY | 1.5605414 | 0.07526882 | 0.09636527 |
| REACTOME_MAPK6_MAPK4_SIGNALING | 1.9018323 | 0.01518438 | 0.01356729 |
| REACTOME_METABOLISM_OF_AMINO_ACIDS_AND_DERIVATIVES | 1.6660275 | 0.02575107 | 0.05610034 |
| REACTOME_METABOLISM_OF_COFACTORS | 1.68201 | 0.01096491 | 0.05198783 |
| REACTOME_METABOLISM_OF_FOLATE_AND_PTERINES | 1.5063604 | 0.06564552 | 0.12242222 |
| REACTOME_METABOLISM_OF_NON_CODING_RNA | 2.1277113 | 0 | 0.00368204 |
| REACTOME_METABOLISM_OF_NUCLEOTIDES | 1.7590241 | 0.00213675 | 0.03215667 |
| REACTOME_METABOLISM_OF_POLYAMINES | 2.0559008 | 0.00214592 | 0.00531063 |
| REACTOME_METABOLISM_OF_PORPHYRINS | 1.3290925 | 0.14401622 | 0.2355269 |
| REACTOME_MICRORNA_MIRNA_BIOGENESIS | 1.8335073 | 0.00441501 | 0.01975593 |
| REACTOME_MISMATCH_REPAIR | 1.692337 | 0.01754386 | 0.04849545 |
| REACTOME_MITOCHONDRIAL_CALCIUM_ION_TRANSPORT | 1.454563 | 0.10183299 | 0.15060389 |
| REACTOME_MITOCHONDRIAL_PROTEIN_IMPORT | 1.9357613 | 0.00222717 | 0.01101033 |
| REACTOME_MITOCHONDRIAL_TRANSLATION | 2.0939116 | 0 | 0.00442696 |
| REACTOME_MITOCHONDRIAL_TRNA_AMINOACYLATION | 1.7093207 | 0.01952278 | 0.04408638 |
| REACTOME_MITOPHAGY | 1.4262942 | 0.14254385 | 0.16855194 |
| REACTOME_MITOTIC_G1_G1_S_PHASES | 1.8995903 | 0.00443459 | 0.01378556 |
| REACTOME_MITOTIC_G2_G2_M_PHASES | 2.0522914 | 0.00448431 | 0.00533021 |
| REACTOME_MITOTIC_METAPHASE_AND_ANAPHASE | 1.9599552 | 0.00223214 | 0.00950266 |
| REACTOME_MITOTIC_SPINDLE_CHECKPOINT | 1.803724 | 0.00881057 | 0.02356927 |
| REACTOME_MRNA_CAPPING | 2.0564995 | 0 | 0.0053212 |
| REACTOME_MRNA_DECAY_BY_3_TO_5_EXORIBONUCLEASE | 1.6946718 | 0.01385681 | 0.04796165 |
| REACTOME_MRNA_SPLICING | 2.203924 | 0 | 0.00192523 |
| REACTOME_MRNA_SPLICING_MINOR_PATHWAY | 2.0965753 | 0 | 0.00457203 |
| REACTOME_MTORC1_MEDIATED_SIGNALLING | 1.6277181 | 0.03950104 | 0.06923267 |
| REACTOME_NEDDYLATION | 1.8047508 | 0.0212766 | 0.02370463 |
| REACTOME_NEGATIVE_EPIGENETIC_REGULATION_OF_RRNA_EXPRESSION | 1.8460368 | 0.00219298 | 0.01855034 |
| REACTOME_NEGATIVE_REGULATION_OF_NOTCH4_SIGNALING | 1.8516599 | 0 | 0.01788137 |
| REACTOME_NEURODEGENERATIVE_DISEASES | 1.5670327 | 0.06540085 | 0.09319289 |
| REACTOME_NONSENSE_MEDIATED_DECAY_NMD | 2.0609376 | 0 | 0.00510986 |
| REACTOME_NONSENSE_MEDIATED_DECAY_NMD_INDEPENDENT_OF_THE_EXON_JUNCTION_COMPLEX_EJC | 1.862448 | 0.00404858 | 0.01676564 |
| REACTOME_NUCLEOBASE_BIOSYNTHESIS | 1.6637809 | 0.03099174 | 0.05662197 |
| REACTOME_NUCLEOTIDE_EXCISION_REPAIR | 2.1506307 | 0 | 0.00346693 |
| REACTOME_NUCLEOTIDE_SALVAGE | 1.7366523 | 0.0021692 | 0.03766616 |
| REACTOME_ORC1_REMOVAL_FROM_CHROMATIN | 1.8660353 | 0 | 0.016304 |
| REACTOME_PCNA_DEPENDENT_LONG_PATCH_BASE_EXCISION_REPAIR | 1.6407242 | 0.04026846 | 0.06466235 |
| REACTOME_PCP_CE_PATHWAY | 1.6494004 | 0.05319149 | 0.06112796 |
| REACTOME_PERK_REGULATES_GENE_EXPRESSION | 1.8863239 | 0.01229508 | 0.01496518 |
| REACTOME_PHOSPHORYLATION_OF_THE_APC_C | 1.7700448 | 0 | 0.02972205 |
| REACTOME_POST_CHAPERONIN_TUBULIN_FOLDING_PATHWAY | 1.5847287 | 0.06420233 | 0.08642687 |
| REACTOME_PREFOLDIN_MEDIATED_TRANSFER_OF_SUBSTRATE_TO_CCT_TRIC | 1.8224194 | 0.00425532 | 0.02150116 |
| REACTOME_PROCESSING_OF_CAPPED_INTRON_CONTAINING_PRE_MRNA | 2.2620168 | 0 | 0.00126037 |
| REACTOME_PROCESSING_OF_CAPPED_INTRONLESS_PRE_MRNA | 1.9051381 | 0.00223714 | 0.01358957 |
| REACTOME_PROCESSIVE_SYNTHESIS_ON_THE_LAGGING_STRAND | 1.5536127 | 0.04072398 | 0.09912106 |
| REACTOME_PROGRAMMED_CELL_DEATH | 1.8827889 | 0.00842105 | 0.01495911 |
| REACTOME_PROTEIN_FOLDING | 1.5005196 | 0.06170213 | 0.12536396 |
| REACTOME_PROTEIN_LOCALIZATION | 1.8311005 | 0.01495727 | 0.02004243 |
| REACTOME_PTEN_REGULATION | 1.8742226 | 0.01301519 | 0.01572336 |
| REACTOME_PURINE_CATABOLISM | 1.5266092 | 0.06485356 | 0.11168794 |
| REACTOME_PYRUVATE_METABOLISM_AND_CITRIC_ACID_TCA_CYCLE | 1.5816333 | 0.06651885 | 0.08791389 |
| REACTOME_RECOGNITION_OF_DNA_DAMAGE_BY_PCNA_CONTAINING_REPLICATION_COMPLEX | 1.7022805 | 0.03363229 | 0.04615493 |
| REACTOME_REGULATION_OF_APOPTOSIS | 1.8427761 | 0 | 0.01865291 |
| REACTOME_REGULATION_OF_EXPRESSION_OF_SLITS_AND_ROBOS | 2.2761066 | 0 | 8.17E-04 |
| REACTOME_REGULATION_OF_MITOTIC_CELL_CYCLE | 1.8712381 | 0 | 0.01584004 |
| REACTOME_REGULATION_OF_MRNA_STABILITY_BY_PROTEINS_THAT_BIND_AU_RICH_ELEMENTS | 2.0325675 | 0 | 0.00534802 |
| REACTOME_REGULATION_OF_PTEN_STABILITY_AND_ACTIVITY | 1.9606483 | 0.00208768 | 0.00970154 |
| REACTOME_REGULATION_OF_RAS_BY_GAPS | 2.0455818 | 0 | 0.00546593 |
| REACTOME_REGULATION_OF_RUNX2_EXPRESSION_AND_ACTIVITY | 1.8783762 | 0.02272727 | 0.0153763 |
| REACTOME_REGULATION_OF_RUNX3_EXPRESSION_AND_ACTIVITY | 2.0407257 | 0 | 0.00544684 |
| REACTOME_REGULATION_OF_TP53_ACTIVITY_THROUGH_PHOSPHORYLATION | 1.8969254 | 0.00454545 | 0.01376046 |
| REACTOME_RESOLUTION_OF_ABASIC_SITES_AP_SITES | 1.7337397 | 0.02073733 | 0.03784657 |
| REACTOME_RESOLUTION_OF_AP_SITES_VIA_THE_MULTIPLE_NUCLEOTIDE_PATCH_REPLACEMENT_PATHWAY | 1.6941433 | 0.02449889 | 0.04796267 |
| REACTOME_RESPIRATORY_ELECTRON_TRANSPORT | 1.960416 | 0.00867679 | 0.00960106 |
| REACTOME_RESPIRATORY_ELECTRON_TRANSPORT_ATP_SYNTHESIS_BY_CHEMIOSMOTIC_COUPLING_AND_HEAT_PRODUCTION_BY_UNCOUPLING_PROTEINS | 1.9164529 | 0.01284797 | 0.01260611 |
| REACTOME_RNA_POLYMERASE_I_PROMOTER_ESCAPE | 1.8431549 | 0.0043384 | 0.01874117 |
| REACTOME_RNA_POLYMERASE_I_TRANSCRIPTION | 1.8778145 | 0.00218341 | 0.01535967 |
| REACTOME_RNA_POLYMERASE_I_TRANSCRIPTION_INITIATION | 2.068652 | 0.00436681 | 0.00497651 |
| REACTOME_RNA_POLYMERASE_I_TRANSCRIPTION_TERMINATION | 1.9953035 | 0.00223214 | 0.00723592 |
| REACTOME_RNA_POLYMERASE_II_PRE_TRANSCRIPTION_EVENTS | 2.3488002 | 0 | 0.00267704 |
| REACTOME_RNA_POLYMERASE_II_TRANSCRIBES_SNRNA_GENES | 2.239864 | 0 | 0.00152622 |
| REACTOME_RNA_POLYMERASE_II_TRANSCRIPTION_ELONGATION | 2.3306353 | 0 | 0.00133852 |
| REACTOME_RNA_POLYMERASE_II_TRANSCRIPTION_PRE_INITIATION_AND_PROMOTER_OPENING | 2.1316564 | 0 | 0.00393958 |
| REACTOME_RNA_POLYMERASE_II_TRANSCRIPTION_TERMINATION | 2.3057413 | 0 | 9.07E-04 |
| REACTOME_RNA_POLYMERASE_III_CHAIN_ELONGATION | 2.016082 | 0 | 0.00609304 |
| REACTOME_RNA_POLYMERASE_III_TRANSCRIPTION | 2.0885127 | 0 | 0.00443711 |
| REACTOME_RNA_POLYMERASE_III_TRANSCRIPTION_INITIATION_FROM_TYPE_1_PROMOTER | 2.0441093 | 0.00234192 | 0.0055129 |
| REACTOME_RNA_POLYMERASE_III_TRANSCRIPTION_INITIATION_FROM_TYPE_3_PROMOTER | 2.1432269 | 0 | 0.00353843 |
| REACTOME_RNA_POLYMERASE_III_TRANSCRIPTION_TERMINATION | 2.0291727 | 0 | 0.00549673 |
| REACTOME_RRNA_MODIFICATION_IN_THE_NUCLEUS_AND_CYTOSOL | 2.0005398 | 0 | 0.00703263 |
| REACTOME_RRNA_PROCESSING | 2.199985 | 0 | 0.00190777 |
| REACTOME_RRNA_PROCESSING_IN_THE_NUCLEUS_AND_CYTOSOL | 2.190382 | 0 | 0.00201543 |
| REACTOME_RUNX1_REGULATES_TRANSCRIPTION_OF_GENES_INVOLVED_IN_DIFFERENTIATION_OF_HSCS | 1.8083436 | 0.01268499 | 0.02341706 |
| REACTOME_S_PHASE | 1.9222745 | 0.0022779 | 0.01219908 |
| REACTOME_SCF_SKP2_MEDIATED_DEGRADATION_OF_P27_P21 | 1.9556371 | 0 | 0.00959038 |
| REACTOME_SELENOAMINO_ACID_METABOLISM | 2.0290227 | 0.00201613 | 0.00541344 |
| REACTOME_SIGNALING_BY_FGFR2_IIIA_TM | 1.8843486 | 0 | 0.01499909 |
| REACTOME_SIGNALING_BY_NOTCH | 1.5436913 | 0.06315789 | 0.10377374 |
| REACTOME_SIGNALING_BY_NOTCH4 | 1.8296386 | 0.01290323 | 0.02023964 |
| REACTOME_SIGNALING_BY_ROBO_RECEPTORS | 2.3095887 | 0.00207469 | 0.00105789 |
| REACTOME_SRP_DEPENDENT_COTRANSLATIONAL_PROTEIN_TARGETING_TO_MEMBRANE | 1.9144826 | 0.00199601 | 0.01278106 |
| REACTOME_STABILIZATION_OF_P53 | 1.9270846 | 0 | 0.01190125 |
| REACTOME_SUMOYLATION_OF_DNA_REPLICATION_PROTEINS | 1.9026752 | 0.00444445 | 0.01366486 |
| REACTOME_SWITCHING_OF_ORIGINS_TO_A_POST_REPLICATIVE_STATE | 1.9500824 | 0 | 0.01009137 |
| REACTOME_SYNTHESIS_OF_ACTIVE_UBIQUITIN:_ROLES_OF_E1_AND_E2_ENZYMES | 1.8224051 | 0 | 0.02137468 |
| REACTOME_TELOMERE_C_STRAND_LAGGING_STRAND_SYNTHESIS | 1.5864693 | 0.04328018 | 0.08613811 |

| | | | |
|---|---|---|---|
| REACTOME_TELOMERE_MAINTENANCE | 1.7377082 | 0.00455581 | 0.03751328 |
| REACTOME_TERMINATION_OF_TRANSLESION_DNA_SYNTHESIS | 1.8017545 | 0.03131991 | 0.02384412 |
| REACTOME_THE_CITRIC_ACID_TCA_CYCLE_AND_RESPIRATORY_ELECTRON_TRANSPORT | 1.9379157 | 0.0155902 | 0.0109947 |
| REACTOME_THE_ROLE_OF_GTSE1_IN_G2_M_PROGRESSION_AFTER_G2_CHECKPOINT | 1.9745617 | 0 | 0.00861197 |
| REACTOME_TNFR2_NON_CANONICAL_NF_KB_PATHWAY | 1.4032557 | 0.12549801 | 0.18304141 |
| REACTOME_TP53_REGULATES_METABOLIC_GENES | 1.686296 | 0.04564316 | 0.05072368 |
| REACTOME_TP53_REGULATES_TRANSCRIPTION_OF_DNA_REPAIR_GENES | 2.2151768 | 0.00225225 | 0.00199453 |
| REACTOME_TRANSCRIPTION_COUPLED_NUCLEOTIDE_EXCISION_REPAIR_TC_NER | 2.0399885 | 0.00455581 | 0.00538842 |
| REACTOME_TRANSCRIPTION_OF_THE_HIV_GENOME | 2.2898886 | 0 | 9.04E-04 |
| REACTOME_TRANSCRIPTIONAL_REGULATION_BY_RUNX1 | 1.6577017 | 0.04237288 | 0.05837716 |
| REACTOME_TRANSCRIPTIONAL_REGULATION_BY_RUNX2 | 1.4940515 | 0.1147541 | 0.1281516 |
| REACTOME_TRANSCRIPTIONAL_REGULATION_BY_RUNX3 | 2.0440857 | 0.00210971 | 0.00541445 |
| REACTOME_TRANSCRIPTIONAL_REGULATION_BY_TP53 | 2.1288748 | 0 | 0.00370308 |
| REACTOME_TRANSLATION | 2.2100782 | 0 | 0.0019223 |
| REACTOME_TRANSLESION_SYNTHESIS_BY_POLH | 1.6726143 | 0.03603604 | 0.05407743 |
| REACTOME_TRANSLESION_SYNTHESIS_BY_POLK | 1.6166819 | 0.06436782 | 0.07381504 |
| REACTOME_TRANSLESION_SYNTHESIS_BY_Y_FAMILY_DNA_POLYMERASES_BYPASSES_LESIONS_ON_DNA_TEMPLATE | 1.8833784 | 0.01342282 | 0.01498709 |
| REACTOME_TRANSPORT_OF_MATURE_TRANSCRIPT_TO_CYTOPLASM | 2.340663 | 0 | 0.0017847 |
| REACTOME_TRISTETRAPROLIN_TTP_ZFP36_BINDS_AND_DESTABILIZES_MRNA | 1.6072316 | 0.04347826 | 0.07738736 |
| REACTOME_TRNA_AMINOACYLATION | 1.8575145 | 0.00223714 | 0.01728679 |
| REACTOME_TRNA_MODIFICATION_IN_THE_NUCLEUS_AND_CYTOSOL | 1.852859 | 0.00442478 | 0.01780637 |
| REACTOME_TRNA_PROCESSING | 2.1589043 | 0 | 0.0033088 |
| REACTOME_TRNA_PROCESSING_IN_THE_NUCLEUS | 2.2465785 | 0 | 0.00153048 |
| REACTOME_UB_SPECIFIC_PROCESSING_PROTEASES | 1.9393744 | 0.0021692 | 0.01089921 |
| REACTOME_UCH_PROTEINASES | 2.1233668 | 0 | 0.00388509 |
| REACTOME_UNFOLDED_PROTEIN_RESPONSE_UPR | 1.4439588 | 0.13179916 | 0.15693074 |
| REACTOME_VIF_MEDIATED_DEGRADATION_OF_APOBEC3G | 1.806462 | 0.00218818 | 0.02347024 |
| REACTOME_VIRAL_MESSENGER_RNA_SYNTHESIS | 2.113393 | 0 | 0.00416071 |
